# A top-down insular cortex circuit crucial for non-nociceptive fear learning

**DOI:** 10.1101/2024.10.14.618356

**Authors:** Junho Han, Boin Suh, Jin-Hee Han

## Abstract

Understanding how threats drive fear memory formation is crucial to understanding how organisms adapt to environments and treat threat-related disorders like PTSD. While traditional Pavlovian conditioning studies have provided valuable insights, the exclusive reliance on electric shock as a threat stimulus has limited our understanding of diverse threats. To address this, we developed a conditioning paradigm using a looming visual stimulus as an unconditioned stimulus (US) in mice and identified a distinct neural circuit for visual threat conditioning. Parabrachial CGRP neurons were necessary for both conditioning and memory retrieval. Upstream neurons in the posterior insular cortex (pIC) responded to looming stimuli, and their projections to the parabrachial nucleus (PBN) induced aversive states and drove conditioning. However, this pIC-to-PBN pathway was not required for foot-shock conditioning. These findings reveal how non-nociceptive visual stimuli can drive aversive states and fear memory formation, expanding our understanding of aversive US processing beyond traditional models.

## Introduction

In dynamic environments, animals must learn to recognize sensory cues and contextual information associated with threats to generate appropriate defensive responses and enhance survival. While adaptive learning is essential, maladaptive fear memories are implicated in anxiety-related disorders, including post-traumatic stress disorder (PTSD). Thus, understanding how threats drive fear memory formation is crucial for developing effective treatments (*1*). To investigate the neural mechanisms of fear learning, Pavlovian fear conditioning has been widely used across various species, particularly rodents (*2*, *3*). In this paradigm, a neutral sensory cue (conditioned stimulus, CS) is paired with an aversive stimulus (unconditioned stimulus, US), leading to fear responses upon CS re-exposure. This model has significantly advanced our understanding of fear memory formation (*4–8*). However, research has predominantly relied on electric foot shocks as the primary US, limiting our understanding of fear learning driven by other types of threats.

In natural environments, threats are not exclusively nociceptive. Predator-associated cues, such as visual or chemical signals, elicit fear-related behaviors like freezing and escape (*9–11*). Although non-nociceptive, these stimuli can induce aversive affective states, leading to fear learning, which is critical for survival by enabling avoidance of physical harm. Previous studies have used live predators and predator odors as non-nociceptive US in rodent fear learning models (*9*, *10*), identifying key brain regions involved in odor-induced fear learning, including the basolateral amygdala (BLA) (*12*), ventral hippocampus (vHPC) (*13*), medial amygdala (MeA) (*14*), and posterior insular cortex (pIC) (*15*). However, findings on odor-based fear conditioning are inconsistent (*16–19*), as different predator odors vary in effectiveness (*10*, *20*) and activate distinct brain regions (*18*). Additionally, odor stimuli are difficult to control precisely, complicating circuit-level investigations.

In contrast, visual threat stimuli offer precise experimental control and cross-species applicability. Rapidly looming visual stimuli have been shown to trigger immediate defensive responses in rodents (*11*, *21*), frogs (*22*), fish (*23*, *24*), primates (*25*), and humans (*26*, *27*). A pioneering study established a visual threat model in mice using an overhead looming dark disc mimicking an approaching aerial predator (*11*). Given their advantages, visual threats seem to offer an ideal framework for developing novel non-nociceptive fear learning models.

Traditionally, US pathways in fear learning have been identified using foot-shock stimuli (*8*, *28–31*). Among these, the thalamus and parabrachial nucleus (PBN) in the brainstem are recognized as key ascending nociceptive pathways. The thalamus has long been considered central to nociceptive US processing due to its widespread projections across the brain (*32–36*). However, recent evidence highlights the PBN as an essential US pathway, transmitting pain-related signals to the central nucleus of the amygdala (CeA) and other downstream regions (*37*). Han et al. demonstrated that calcitonin gene-related peptide (CGRP)-expressing neurons in the PBN (CGRP^PBN^ neurons) are essential for foot-shock conditioning in mice and that optogenetic activation of CGRP^PBN^-CeA pathway induces fear learning (*37*).

Interestingly, CGRP^PBN^ neurons respond not only to electric shocks but also to various non-nociceptive stimuli (*38*, *39*), including visual threats, suggesting a potential role in non-nociceptive fear learning. However, whether CGRP^PBN^ neurons process US signals during non-nociceptive fear learning remains unclear. Furthermore, unlike nociceptive pathways, which are known to originate from the spinal cord and periaqueductal gray (PAG) (*40–42*), the upstream sources conveying non-nociceptive input to PBN neurons remain unidentified.

Here, we developed a threat conditioning paradigm in mice using the previously established looming visual threat. Through this model, we identified PBN^CGRP^ neurons and a distinct pIC-to-PBN top-down circuit as essential for visual threat conditioning. Notably, this circuit was not required for foot-shock conditioning. Furthermore, the pIC-to-PBN circuit was sufficient to induce aversive affective states and drive threat conditioning as a US. These findings highlight its role in processing threat-relevant, non-nociceptive visual information, independent of sensory pain pathways.

## Results

### Looming Visual Stimuli Drive Threat Conditioning as a US

We aimed to develop a visual threat conditioning model using a looming visual stimulus as a US. The stimulus consisted of a rapidly expanding dark disc (5° to 35° visual angle in 300 ms), which remained at full size for 500 ms. First, we confirmed that this stimulus induces innate defensive responses in mice. Exposure to 10 consecutive looming stimuli elicited robust freezing, either immediately or following an escape response (**Fig. 1, A to C**). Eight animals exhibited immediate freezing, while the remaining two initially escaped before freezing. A previous study (*43*) reported predominant escape-freezing responses over immediate freezing in a no-shelter condition. However, differences in experimental conditions, particularly looming stimulus parameters, may account for these discrepancies. Notably, another study (*44*) used looming stimuli similar to ours and reported that mice primarily exhibited immediate freezing in a no-shelter condition, consistent with our findings.

**Fig. 1.**
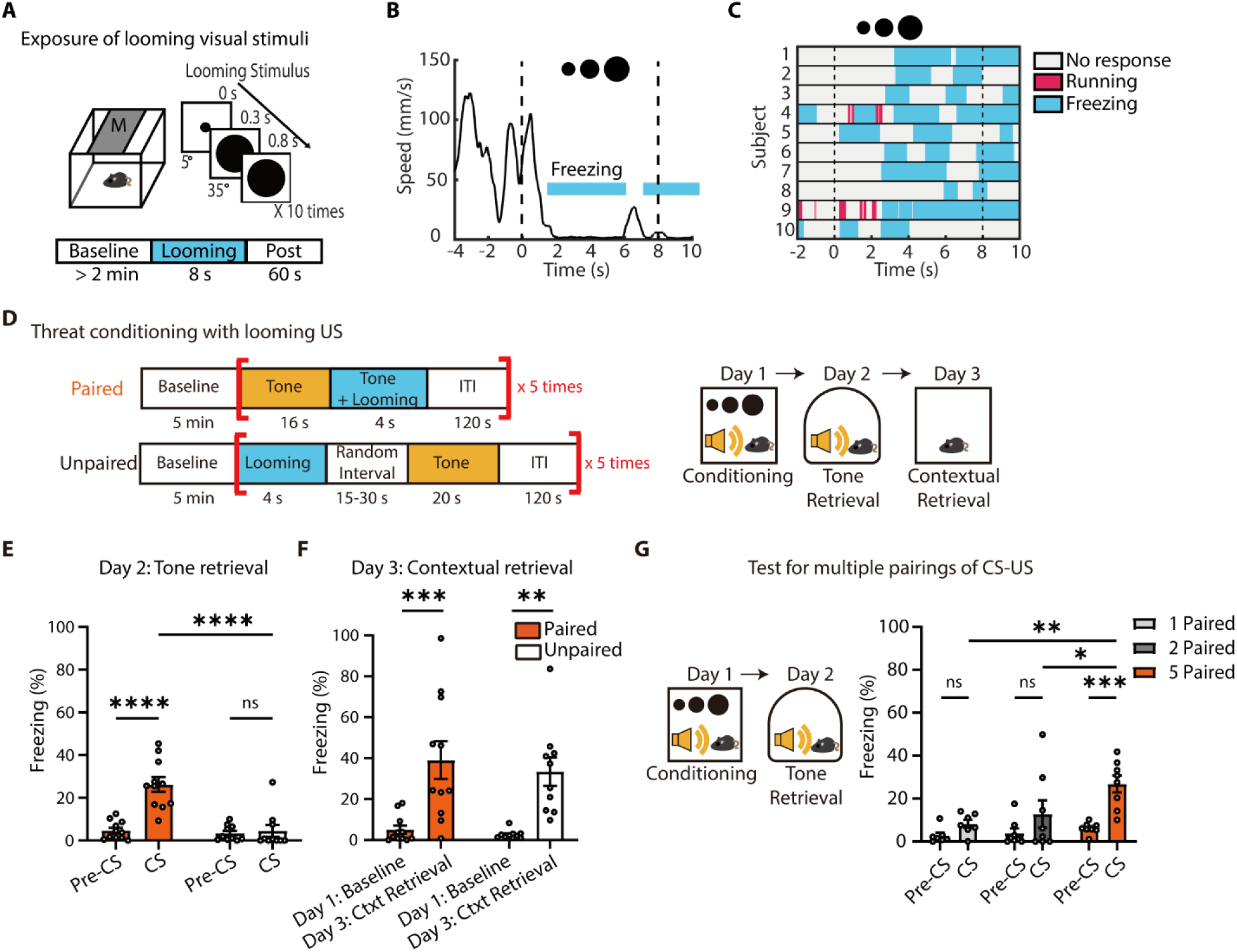
Looming visual stimuli drive formation of auditory and contextual fear memory in mice. (**A**) Schematic of behavioral tests for exposure of looming visual stimuli (a 10-times repeat of a single looming for 8 s). (**B**) Changes in movement speed of mice in response to looming visual stimuli. The blue shaded bars indicate a period during which mice displayed freezing behavior. (**C**) Ethogram showing defensive behavioral responses to the presentation of looming visual stimuli. Mice displayed robust freezing, either immediately or following an escape response (*N*= 10, 8 mice showed immediate freezing and other 2 mice showed escape-freezing). (**D**) Schematic of a threat conditioning protocol with looming visual stimuli. (**E**) Auditory fear memory was formed only in the paired group (Paired group, *N*= 11; Unpaired group, *N*= 10; RM two-way ANOVA, Time × Group, *F*_1,19_= 17.2, *P*= 0.0005). (**F**) Both paired and unpaired groups show elevated levels of contextual freezing during re-exposure to the conditioning chamber on day3 (RM two-way ANOVA, Time, *F*_1,19_= 36.38, *P*< 0.0001). (**G**) Single and double pairings of tone CS and looming US were not effective in driving the formation of auditory fear memory (Single-paired group, *N*= 7; Double-paired group, *N*= 8; Five-times paired group, *N*= 8; RM two-way ANOVA, Group, *F*_2,20_= 4.500, *P*= 0.0243; right). All data are mean ± s.e.m. *p < 0.05. **p < 0.01. ***p < 0.001. ****p < 0.0001. ns, not significant.

Given that looming stimuli pose no actual harm, we tested for habituation. Mice exposed to looming stimuli across five days showed no significant reduction in freezing (**fig. S1A, B**). However, when the same stimulus was repeatedly presented within short intervals (five stimuli per session, repeated 10 times with 120-s intertrial intervals [ITI]), freezing responses declined after the 7th exposure, indicating habituation (**fig. S1D, E**). This reduction in freezing was not due to a shift in defensive behavioral patterns, as repeated looming exposures did not increase escape behavior probability (**fig. S1C, F**). While significant habituation was observed after the 7th trial, some mice exhibited a decrease in freezing levels to looming stimuli relative to baseline freezing starting from the 6th trial (**fig. S1G**). Based on these findings, we used five CS-US pairings in subsequent conditioning experiments.

To determine whether looming stimuli can serve as an effective US, we paired a neutral auditory tone (2.7 kHz, 80 dB) as a CS with five consecutive looming stimuli as a US (looming US). In the paired group, the tone was presented for 20 s, overlapping with the final 4 s of the looming US. In the unpaired group, the US and CS were presented separately with a randomized 15-to 30-s interval (**Fig. 1D**). During the retrieval test in a different context, only the paired group exhibited significant freezing to the tone (**Fig. 1E**). Contextual fear memory was also assessed by re-exposing mice to the original conditioning chamber, where both groups showed significant freezing (**Fig. 1F**). Additional experiments revealed that fewer than five CS-US pairings failed to establish robust conditioning (**Fig. 1G**). These results demonstrate that a non-nociceptive visual threat can drive robust fear learning as a US in mice.

### CGRP^PBN^ neurons selectively respond to looming US and acquired tone CS, but not to neutral stimuli

The PBN is a major ascending nociceptive pathway with extensive connections to the CeA and hypothalamus (*45*, *46*), regulating emotional behaviors and autonomic responses. This suggests its primary role in the affective pain processing rather than sensory pain perception. Affective pain, defined as the emotional unpleasantness of pain (*47–50*), drives avoidance behavior, facilitating fear learning. Supporting this, CGRP^PBN^ neurons are essential for foot-shock conditioning, and optogenetic activation of the CGRP^PBN^-CeA pathway induces fear memory formation (*37*). Beyond nociceptive processing, CGRP^PBN^ neurons also respond to various non-nociceptive aversive stimuli (*39*) and regulate food neophobia (*38*), suggesting a broader function in aversive state encoding. These findings indicate that the PBN serves as a general danger-detection hub (*51*), integrating diverse aversive signals and regulating emotional behaviors. Thus, we reasoned that CGRP^PBN^ neurons may also contribute to fear memory formation in response to visual threats.

First, to verify whether CGRP^PBN^ neurons specifically respond to looming visual stimuli, we performed in vivo fiber photometry calcium recordings in CGRP^PBN^ neurons during exposure to looming and flickering stimuli (a non-threatening control; **Fig. 2A**). Cre-dependent AAV encoding GCaMP6m was injected into the PBN of *Calca^Cre^* transgenic mice (*52*), in which Cre recombinase is specifically expressed in CGRP+ neurons, with an optic fiber implanted above the injection site (**Fig. 2B**). While flickering stimuli failed to induce freezing, looming threats elicited strong freezing responses (**Fig. 2C**). Correspondingly, CGRP^PBN^ neurons exhibited no calcium activation in response to flickering stimuli but showed significant and sustained activation (∼1 min) following looming threats (**Fig. 2, D to G**). These results demonstrate that CGRP^PBN^ neurons selectively respond to threatening visual stimuli rather than neutral visual cues.

**Fig. 2.**
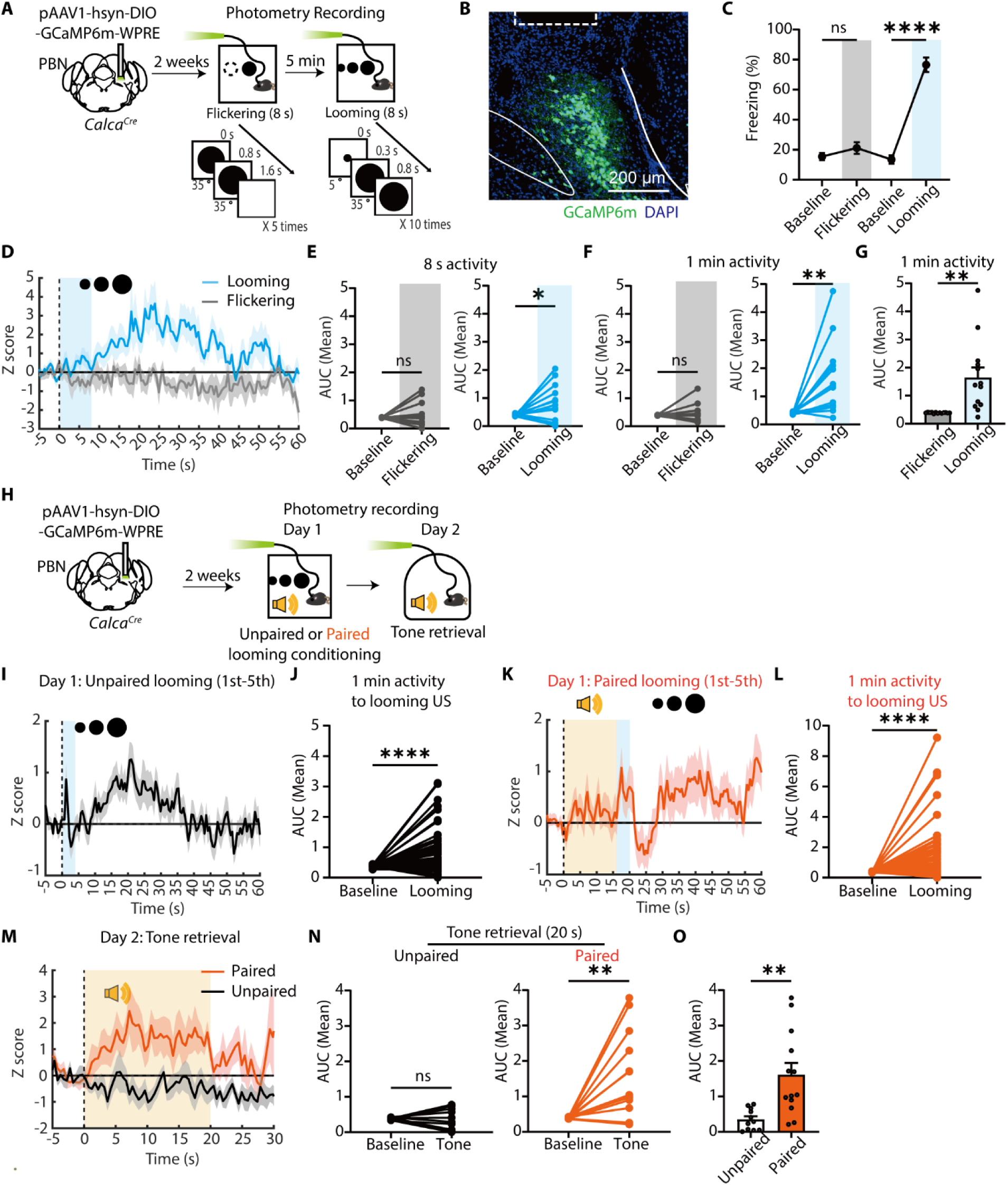
CGRP^PBN^ neurons selectively respond to looming US and acquired tone CS, but not to neutral stimuli. (**A**) Schematic of fiber photometry recordings in CGRP^PBN^ neurons during visual stimuli. (**B**) Representative image of GCaMP6m expression in the PBN; the dotted rectangle marks the optic fiber location. (**C**) Freezing levels during flickering and looming stimuli (*N*= 13 mice; RM one-way ANOVA, Time, *F*_12,36_= 2.786, *P*= 0.0087). (**D)** Z-scored calcium activity (mean ± s.e.m.) during visual stimuli. (**E, F**) Average area under the curve (AUC) comparison of calcium activity during visual stimuli (E; flickering; *N*= 13 mice; two-tailed paired t-test, t_12_= 0.6077, *P*= 0.5547; looming; t_12_= 2.221, *P*= 0.0464) and during 1 min (F; flickering; t_12_= 0.6251, *P*= 0.5436; looming; t_12_= 3.482, *P*= 0.0045). (**G**) AUC comparison between flickering and looming stimuli (t_12_= 3.541, *P*= 0.0041). (**H**) Schematic of fiber photometry recordings of CGRP^PBN^ neurons during visual threat conditioning. (**I-J**) Z-scored calcium activity of CGRP^PBN^ neurons in the unpaired group during the 1st-5th exposures to looming US (J; *N*= 55 trials; two-tailed Wilcoxon matched-pairs signed rank test, W= −1042, *P*< 0.0001). (**K-L**) Z-scored calcium activity of CGRP^PBN^ neurons in the paired group during the 1st-5th exposures to looming US (L; N = 65 trials; two-tailed Wilcoxon matched-pairs signed-rank test, W = −1261, *P* < 0.0001). (**M**) Z-scored calcium activity of CGRP^PBN^ neurons during the tone retrieval test. (**N**) Average AUC of calcium activity in the unpaired (left; two-tailed paired t-test, t_10_= 0.3602, *P*= 0.7262) and paired groups (right; t_12_= 3.714, *P*= 0.0030) during the tone retrieval test. (**O**) Comparison of AUC between the two groups during the tone retrieval test (Two-tailed unpaired t-test, t_22_= 3.431, *P*= 0.0024). All data are mean ± s.e.m. *p < 0.05. **p < 0.01. ****p < 0.0001. ns, not significant.

Next, we examined CGRP^PBN^ activity during visual threat conditioning and tone retrieval tests. Mice were divided into paired and unpaired conditioning groups (**Fig. 2H**). As expected, both groups showed increased CGRP^PBN^ activity in response to the looming US during conditioning (**Fig. 2, I to L**). However, during retrieval, only the paired group exhibited increased calcium activity in response to the tone CS (**Fig. 2, M to O**). These findings suggest that following conditioning, CGRP^PBN^ neurons encode the acquired danger value of the tone CS, likely through synaptic plasticity-dependent mechanisms.

### CGRP^PBN^ neurons are required for fear memory formation and its retrieval, but not for innate defensive response to a visual threat

To assess the necessity of CGRP^PBN^ neurons in visual threat conditioning, we inactivated them using tetanus toxin light chain (TetTox)(*53*). Cre-dependent AAV encoding TetTox and dTomato was injected into the PBN of *Calca^Cre^* mice, with tdTomato alone serving as a control (**Fig. 3A**). Post hoc histological analysis confirmed restricted viral expression in the PBN (**Fig. 3B**). Open-field tests confirmed no locomotor or anxiety differences between groups before conditioning (**Fig. 3C**).

**Fig. 3.**
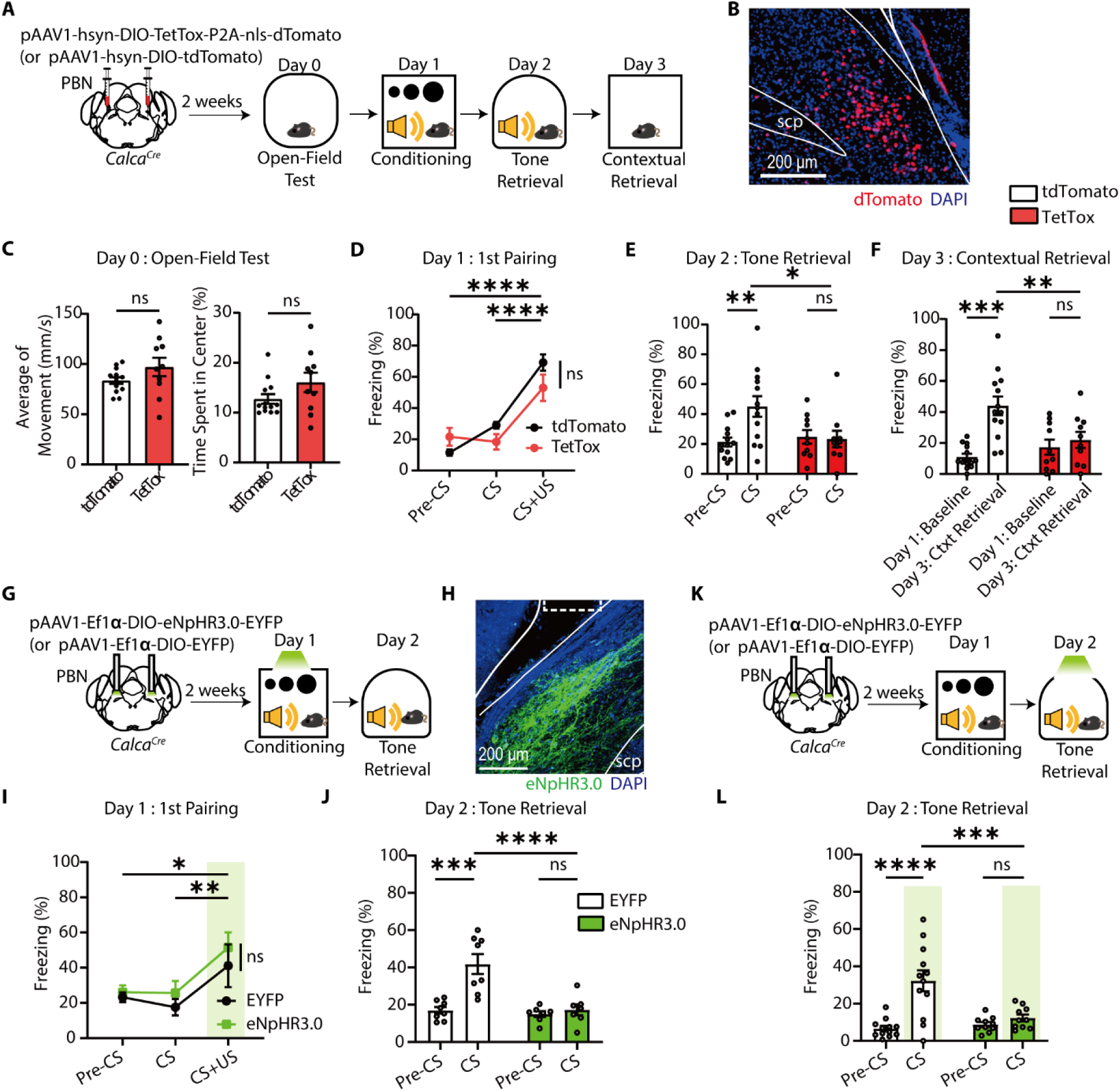
The inhibition of CGRP^PBN^ neurons impairs fear memory formation and retrieval but not innate defensive behavior to looming stimuli. (**A**) Schematic of TetTox-mediated inhibition of CGRP^PBN^ neurons during visual threat conditioning. (**B**) Representative image showing the expression of dTomato in the PBN, especially in the external lateral subdivision of PBN (PBel). Scale bars, 200 µm. SCP, superior cerebellar peduncle. (**C**) Average of movement (left; TetTox, *N*= 10; tdTomato, *N* = 13; two-tailed unpaired *t*-test, *t*_21_= 1.518, *P*= 0.1439) and center zone time (right, two-tailed Mann Whitney test, U= 46, *P*= 0.2569) were comparable between groups. (**D**) Freezing levels during the first CS–US pairing on Day 1 (RM two-way ANOVA, Time × Group, *F*_2,42_= 5.334, *P*= 0.0086). (**E-F**) TetTox inhibition of CGRP^PBN^ neurons impairs auditory (E; RM two-way ANOVA, Time × Group, *F*_1,21_= 7.251, *P*= 0.0136) and contextual fear memory (F; RM two-way ANOVA, Time × Group, *F*_1,21_= 7.798, *P*= 0.0109). (**G**) Schematic of optogenetic inhibition of CGRP^PBN^ neurons during visual threat conditioning. (**H**) Representative image of eNpHR3.0-EYFP expression in CGRP^PBN^ neurons; the dotted rectangle marks the optic fiber tip location. Scale bars, 200 µm. (**I**) Freezing during the first CS–US pairing (eNpHR3.0, *N*= 7; EYFP, *N*= 8; RM two-way ANOVA, Time, *F*_2,26_= 7.167, *P*= 0.0033). (**J**) Inhibition of CGRP^PBN^ neurons during visual US conditioning impaired the formation of auditory fear memory (RM two-way ANOVA, Time × Group, *F*_1,13_= 9.965, *P*= 0.0076). (**K**) Schematic of optogenetic inhibition of CGRP^PBN^ neurons during auditory memory retrieval. (**L**) Inhibition of CGRP^PBN^ neurons impaired retrieval of auditory fear memory (eNpHR3.0, *N*= 10; EYFP, *N*= 12; RM two-way ANOVA, Time × Group, *F*_1,20_= 18.16, *P*= 0.0004). All data are mean ± s.e.m. *p < 0.05. **p < 0.01. ***p < 0.001. ****p < 0.0001. ns, not significant.

During conditioning, both TetTox and control groups exhibited similar freezing to the looming US (first CS-US pairing), indicating that CGRP^PBN^ neurons are not essential for innate defensive responses to visual threats (**Fig. 3D**), consistent with prior studies (*39*). This finding was further validated in an independent experiment (**fig. S2, A and B**). However, fear memory formation was significantly impaired—TetTox-expressing mice showed no significant increase in freezing to the conditioned tone or context, whereas control mice exhibited robust freezing (**Fig. 3, E and F**). This suggests that CGRP^PBN^ neurons are required for fear learning but not for reflexive defensive behaviors to looming stimuli.

To further investigate the role of CGRP^PBN^ neurons in visual threat conditioning and memory retrieval, we performed optogenetic inhibition. Cre-dependent AAV encoding eNpHR3.0-EYFP or EYFP alone was injected into the PBN of *Calca^Cre^* mice, with an optic fiber implanted above the injection site. Light delivery inactivated CGRP^PBN^ neurons throughout conditioning, beginning at the onset of the first US presentation and continuing until the session concluded (**Fig. 3G**). Histological analysis confirmed highly restricted eNpHR3.0 expression in the PBN (**Fig. 3H**).

Consistent with the TetTox experiment, immediate freezing responses (measured during first CS-US pairing) were unaffected (**Fig. 3I**). However, fear memory formation was abolished in the eNpHR3.0 group (**Fig. 3J**). In a separate experiment, inhibition of CGRP^PBN^ neurons during retrieval, rather than conditioning, resulted in significantly impaired freezing to the tone CS, indicating a role in memory retrieval (**Fig. 3, K and L**). Together, these findings demonstrate that CGRP^PBN^ neurons are required for fear memory formation and retrieval but are dispensable for innate defensive responses to visual threats.

### pIC and SC cells projecting to the CGRP^PBN^ neurons are activated by looming visual stimuli

CGRP^PBN^ neurons are critical for visual threat conditioning, but their upstream non-nociceptive inputs remain unclear. While nociceptive stimuli are relayed via the spinal cord or PAG (*37*, *40–42*), the pathway conveying non-nociceptive US information is less understood. To identify CGRP^PBN^ inputs, we performed monosynaptic retrograde tracing by injecting Cre-dependent AAV-hsyn-DIO-TVA-HA-N2cG into the PBN of *Calca^Cre^*mice, followed by SADB19-GFP rabies virus (**fig. S3A**). Starter cells were localized exclusively in the PBN (**fig. S3B**), and retrogradely labeled neurons were observed in ipsilateral and contralateral brain regions associated with emotional processing, sensory integration, and motor functions, including the pIC, CeA, PAG, and superior colliculus (SC) (**fig. S3, C to E**).

To determine which of these inputs are activated by visual threats, we analyzed c-Fos expression in projection neurons following looming or flickering stimuli. Mice injected bilaterally with AAV helper and rabies virus were exposed to either looming or flickering stimuli (**Fig. 4A, fig. S4A**). Only looming stimuli significantly increased freezing behavior (**Fig. 4B**). We analyzed c-Fos expression in fear- and vision-related upstream regions, including the insular cortex (IC), CeA, PAG, and SC. Anatomically, CGRP^PBN^-projecting neurons were identified in both the anterior (aIC) and posterior IC (pIC) in layers 5 and 6 (**fig. S5, B and C**), but projections from the pIC were notably denser (**fig. S5D**). Given that the pIC is more closely associated with pain and aversive states, whereas the aIC is primarily linked to appetitive processing (*54–56*), we focused on the pIC as a key candidate for relaying aversive affective signals. In the pIC, most GFP+ cells showed significantly greater activation in the looming group than in the flickering group (**Fig. 4C, D, and fig. S5A, B**). Similarly, SC GFP+ neurons in the intermediate layers exhibited significantly increased activation to looming (**Fig. 4, C, G**). This pIC and SC-specific activation was restricted to GFP+ projection neurons, as no group differences were found in GFP-populations across brain regions (**Fig. 4, C to G**). In contrast, no activation differences were observed in the CeA or PAG (**Fig. 4, C, E, and F**). These findings suggest that projection neurons in the pIC (pIC^→CGRP-PBN^) and SC are selectively activated by looming threats and may serve as upstream sources of US information for CGRP^PBN^ neurons.

**Fig. 4.**
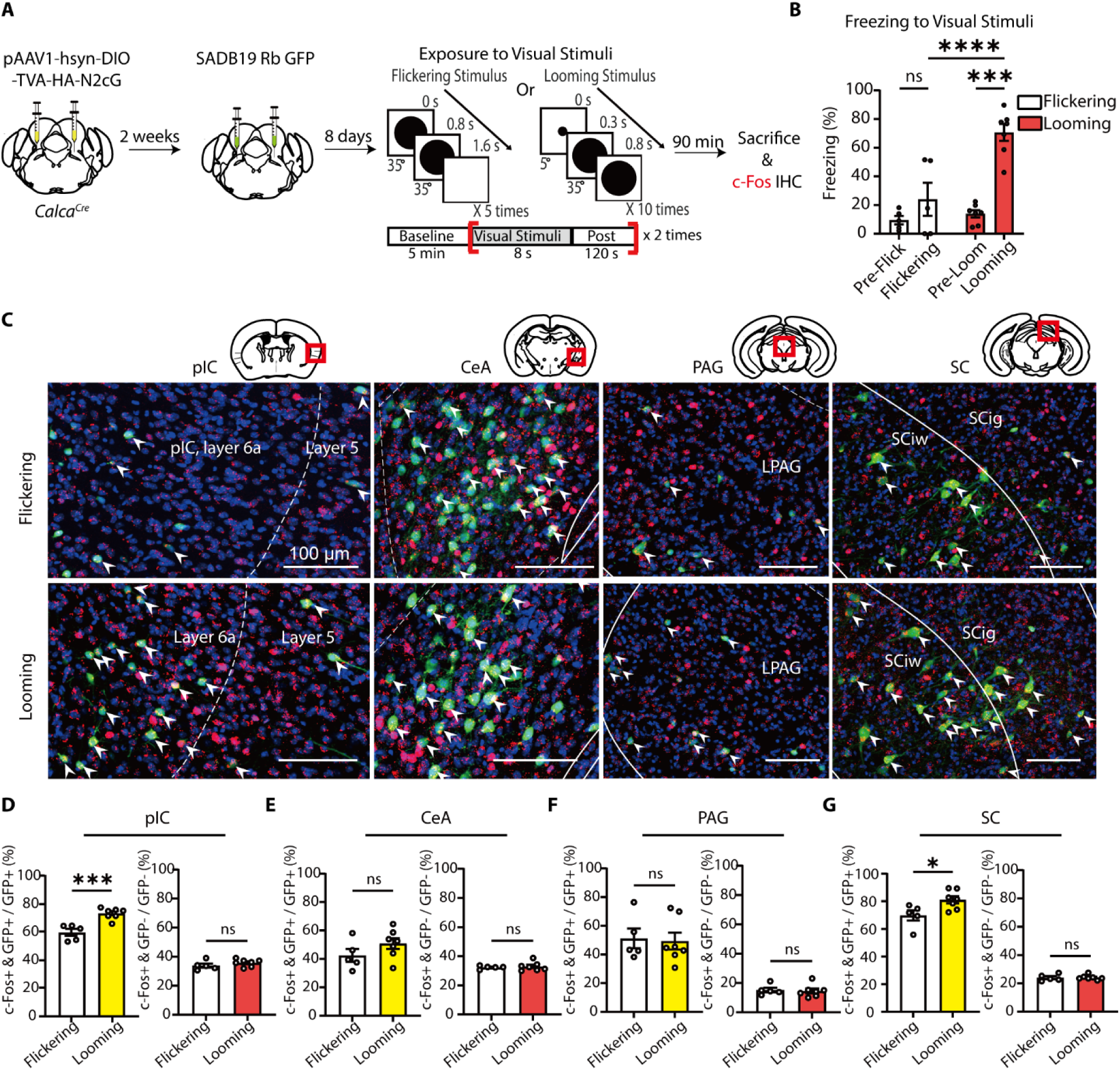
Identification of CGRP^PBN^-projecting neurons activated by looming visual stimuli. (**A**) Schematic of a monosynaptic retrograde tracing of CGRP^PBN^ neurons and c-Fos imaging in response to looming stimuli. Bilateral injections of AAV2/1-hsyn-DIO-TVA-HA-N2cG into the PBN of *Calca^Cre^* mice were followed by bilateral injections of SADB19-GFP rabies virus into the same site. (**B**) Looming visual stimuli induced a robust freezing response in mice, whereas flickering stimuli did not (Flickering, *N*= 5, Looming, *N*= 7; RM two-way ANOVA, Time × Group, *F*_1,10_= 9.893, *P*= 0.0104). (**C**) Representative confocal microscopic images showing CGRP^PBN^-projecting GFP+ neurons (Green) and c-Fos+ neurons (Red) in the pIC, CeA, PAG, and SC. A white arrowhead indicates co-localized neurons. Images of starter cells in the PBN are presented in the fig. S4. Scale bars, 100 µm. LPAG, lateral periaqueductal gray; SCiw, intermediate white layer of the superior colliculus; SCig, intermediate gray layer of the superior colliculus. (**D-G**) The ratio of double-positive neurons (GFP+ & c-Fos+) within GFP+ or GFP- cell populations was compared between the groups (Flickering vs Looming) in each analyzed region: pIC (D) (Flickering, *N*= 5; Looming, *N*= 7; left; Two-tailed unpaired *t*-test, *t*_10_= 4.952, *P*= 0.0006; right; *t*_10_= 1.145, *P*= 0.2787), CeA (E) (left; *t*_10_= 1.404, *P* = 0.1906; right; *t*_10_= 0.4580, *P*= 0.6567), PAG (F) (left; *t*_10_= 0.2072, *P* = 0.8400; right; *t*_10_= 0.2809, *P*= 0.7845), and SC (G) (left; *t*_10_= 2.593, *P*= 0.0268; right; *t*_10_= 0.1301, *P*= 0.8901). All data are mean ± s.e.m. *p < 0.05. ***p < 0.001. ****p < 0.0001. ns, not significant.

### Selective activation of pIC^→CGRP-PBN^ neurons in response to threat stimuli

The IC is well known for processing negative emotions and contributing to fear, anxiety, and disgust (*48*, *57*). In humans, IC activity correlates with affective and social pain, and lesions can lead to pain asymbolia, where pain is perceived without emotional distress (*48*, *58–60*). In rodents, optogenetic activation of the pIC induces an aversive state (*56*), and pIC activity is required for maintaining learned fear (*61*), suggesting its role in encoding aversive affect rather than sensory pain.

The IC has also been implicated in non-nociceptive fear conditioning. In humans, exposure to aversive images as a US increases IC activation in response to both the US and CS (*62*, *63*). In rodents, inactivation of the pIC impairs predator odor fear conditioning and memory recall (*15*). Notably, the IC has extensive reciprocal connections with the PBN (*45*, *46*), and we identified the increased activity of pIC^→CGRP-PBN^ neurons in response to looming stimuli (**Fig. 4D**). Based on these findings, we hypothesized that the pIC transmits threat-relevant aversive affective signals to the CGRP^PBN^ neurons to drive visual threat conditioning.

To further prove whether pIC^→CGRP-PBN^ neurons selectively respond to threat stimuli, we performed photometry calcium recordings. Using a rabies-virus-based Flp system, we expressed GCaMP6s selectively in pIC^→CGRP-PBN^ neurons (**Fig. 5, A–C**). Mice were exposed to either looming and foot shock stimuli or flickering and foot shock stimuli. Photometry recordings revealed that pIC^→CGRP-PBN^ neurons were persistently activated by both looming and foot shock stimuli but exhibited distinct activation dynamics: a slow increase to looming and a rapid increase to foot shock (**Fig. 5, D–F**). In contrast, flickering stimuli did not induce pIC^→CGRP-PBN^ activation, whereas foot shock elicited a robust response (**Fig. 5, G–I**). These findings indicate that pIC^→CGRP-PBN^ neurons selectively respond to threats rather than general sensory stimuli.

**Fig. 5.**
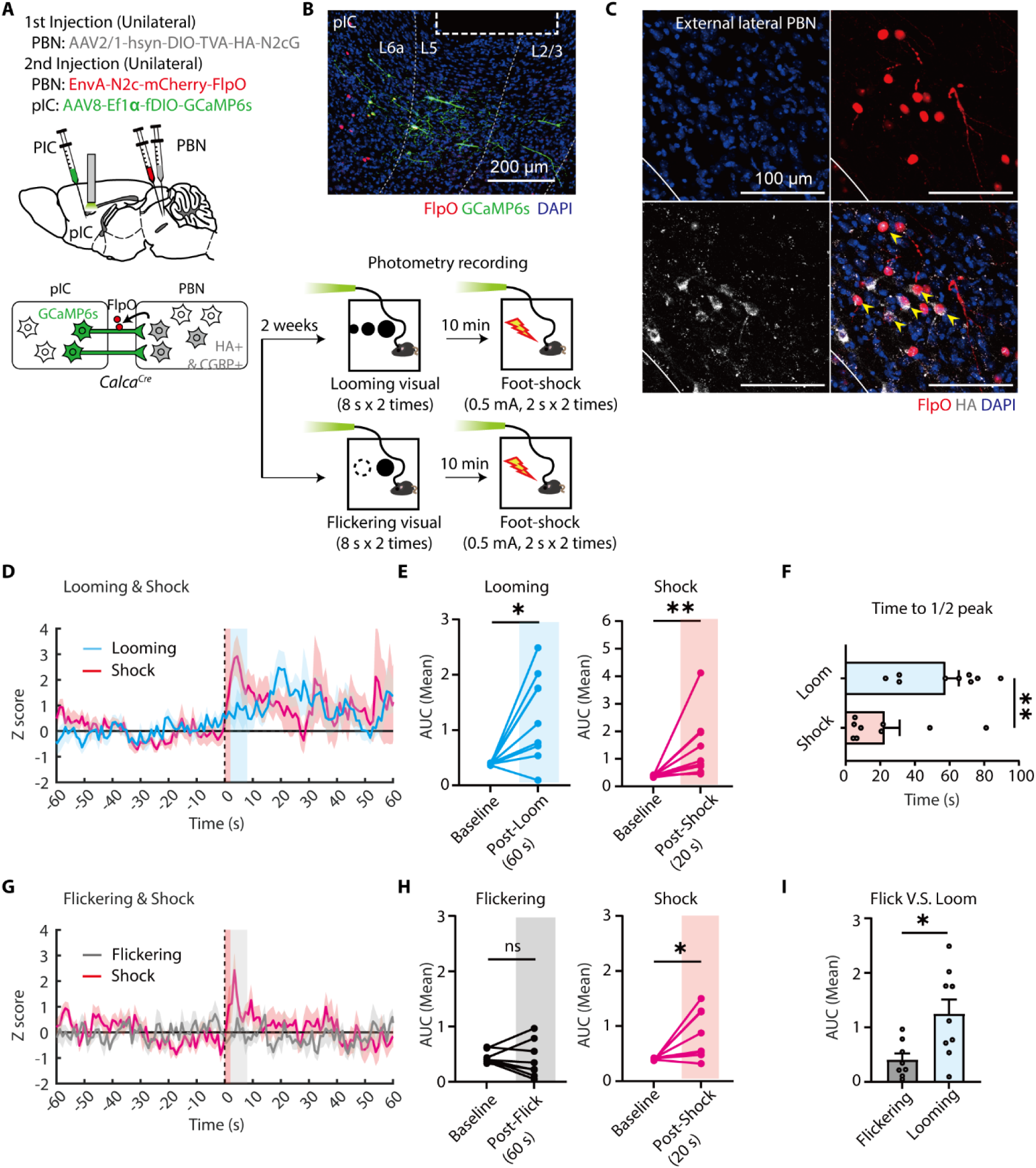
Fiber photometry recordings of pIC^→CGRP-PBN^ neuronal activity in response to visual and foot-shock stimuli. **(A)** Schematic of fiber photometry recordings of pIC^→CGRP-PBN^ neuronal responses to looming (or flickering) & foot-shock stimuli. **(B)** Representative image of GCaMP6s (green) and FlpO (red) expression in the pIC. All GCaMP6s+ cells were FlpO+. The dotted rectangle marks the optic fiber location. Scale bar, 200 µm. **(C)** Representative images of starter cells in the PBN. Yellow arrowheads indicate FlpO-mCherry+ & HA+ starter cells. Scale bars, 100 µm. **(D)** Z-scored calcium activity (mean ± s.e.m) in response to looming and foot-shock stimuli (the blue shaded area: 8-s looming stimuli; the red shaded area: 2-s foot-shock). **(E)** Average during looming and foot-shock (*N*= 9 mice; left; baseline = −60 to 0 s, looming = 0-60 s after the onset of visual stimuli; Two-tailed paired t-test, t_8_= 3.291, *P*= 0.011; right; shock = 0-20 s after the onset of shock stimuli; Two-tailed Wilcoxon matched-pairs signed rank test, W= 45, *P*= 0.0039). **(F)** Time to reach half of the peak z-score after the peak (Two-tailed Mann Whitney test, U= 11, *P*= 0.0072). **(G)** Z-scored calcium activity (mean ± s.e.m) during flickering and foot-shock (the gray shaded area: 8-s flickering stimuli; the red shaded area: 2-s foot-shock). **(H)** Average AUC during flickering and foot-shock (*N*= 8 mice; left; baseline = −60 to 0 s, flickering = 0-60 s after the onset of visual stimuli; Two-tailed Wilcoxon matched-pairs signed rank test, W= −6, *P*= 0.7422; right; shock = 0-20 s after the onset of shock stimuli; Two-tailed paired t-test, t_7_= 2.791, *P*= 0.0269). **(I)** Comparison of AUC between flickering and looming (Two-tailed unpaired t-test, t_15_= 2.798, *P*= 0.0135). All data are presented as mean ± s.e.m. *p < 0.05, **p < 0.01; ns, not significant.

Moreover, the sustained activity of pIC^→CGRP-PBN^ neurons beyond threat exposure suggests they encode persistent aversive states rather than transient threat detection (*56*, *64*, *65*). Their prolonged response to looming stimuli also parallels the sustained activation observed in CGRP^PBN^ neurons (**Fig. 2D**), suggesting that pIC input may contribute to prolonged CGRP^PBN^ activation.

### pIC^→PBN^ neurons are required for CGRP^PBN^ neuronal activation by visual threats but not to foot shock

If the pIC transmits threat-relevant signals to CGRP^PBN^ neurons, then its input should be required for CGRP^PBN^ activation. To test this, we inhibited pIC neurons projecting to the PBN (pIC^→PBN^) using DREADDs (Designer Receptors Exclusively Activated by Designer Drugs) while monitoring CGRP^PBN^ activity in response to looming and foot shock. Cre-dependent AAV-hsyn-DIO-GCaMP6m was injected into the PBN of *Calca^Cre^*mice, and an optic fiber was implanted above the injection site (**Fig. 6A**). To selectively express hM4Di DREADDs in pIC^→PBN^ neurons, AAVretro-FlpO was injected into the PBN, followed by Flp-dependent AAV encoding hM4Di-mCherry (or mCherry as a control) into the pIC (**Fig. 6B**).

**Fig. 6.**
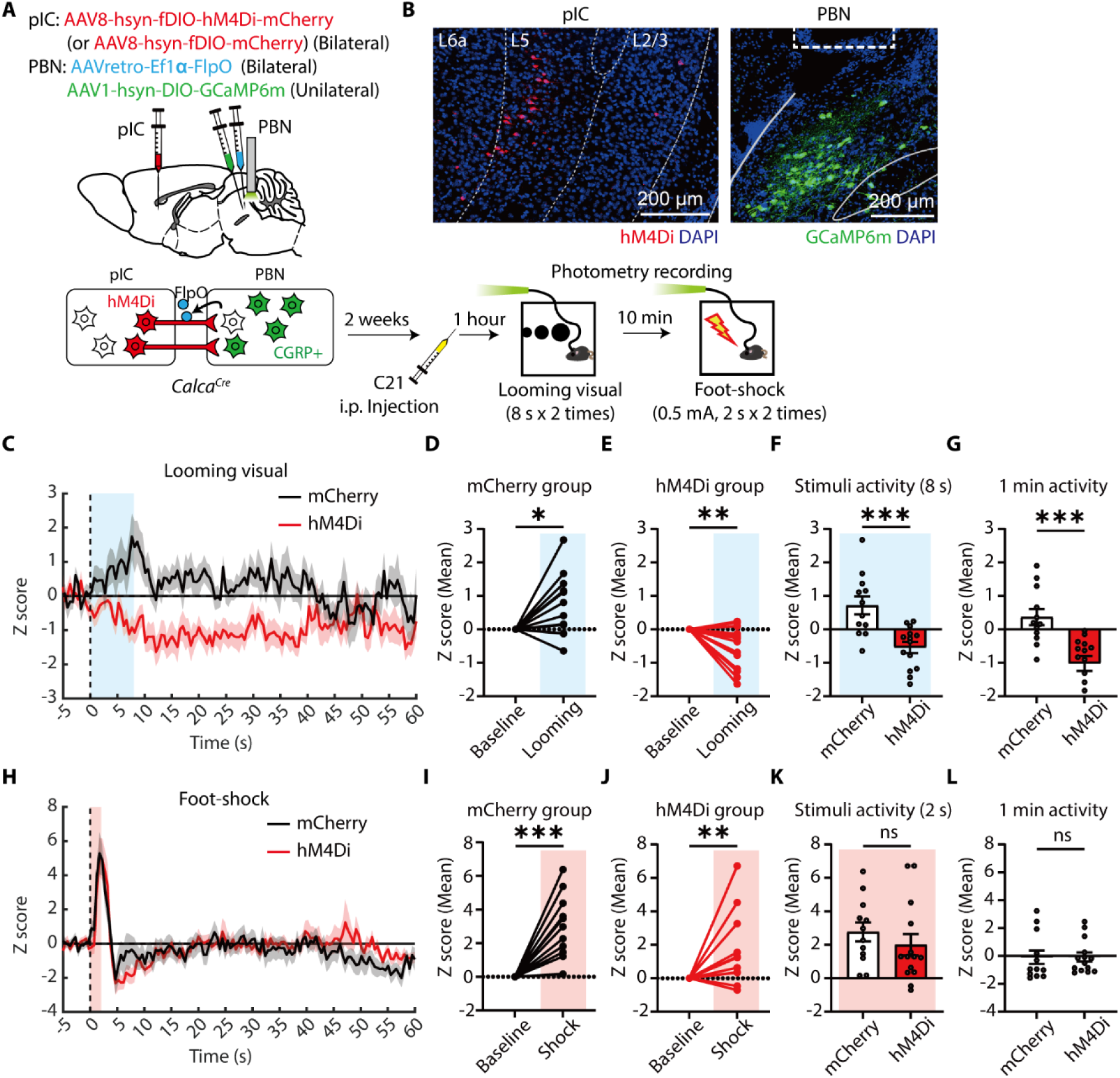
pIC^→PBN^ neurons are necessary for activation of CGRP^PBN^ neurons to looming but not foot-shock stimuli. (**A**) Schematic of photometry recording of CGRP^PBN^ neurons with hM4Di inhibition of pIC^→PBN^ neurons. (**B**) Representative images showing expression of hM4Di-mCherry in the pIC^→PBN^ neurons (left) and GCaMP6m in CGRP^PBN^ neurons (right). The dotted rectangle indicates the location of optic fiber. Scale bar, 200 µm. (**C**) Z-scored calcium activities of CGRP^PBN^ neurons to visual stimuli (blue shaded area: 8-s looming stimuli). (**D-E**) Average calcium activity during visual stimuli in the mCherry group (D) (baseline = −5 to 0 s, looming = 0 to 8 s; *N*=12 trials; 2 trials per mouse, 6 mice; one sample t-test, t_11_= 2.659, *P*= 0.0222) and hM4Di group (E) (*N* =14 trials; 2 trials per mouse, 7 mice; t_13_= 3.266, *P*= 0.0061). (**F-G**) Comparison of average changes in calcium activity evoked by looming stimuli between two groups during stimuli (F) (Two-tailed unpaired t-test, t_24_= 4.104, *P*= 0.0004) and for 1min (G) (Two-tailed unpaired t-test, t_24_= 4.203, *P*= 0.0003). (**H**) Z-scored calcium activities of CGRP^PBN^ neurons to foot-shock stimuli (red shaded area: 2-s foot-shock). (**I-J**) Average calcium activity during foot-shock in the mCherry group (I) (baseline= −5 to 0 s, shock= 0 to 2 s; *N* =12 trials; one sample t-test, t_11_= 4.887, *P*= 0.0005) and hM4Di group (J) (*N* =14 trials; one sample Wilcoxon test, W= 89, *P*= 0.0031). (**K-L**) Comparison of average changes in calcium activity evoked by foot-shock stimuli between two groups during stimuli (K) (Two-tailed Mann Whitney test, U= 63, *P*= 0.2972) and for 1 min (L) (Two-tailed Mann Whitney test, U= 69, *P*= 0.4624). All data are mean ± s.e.m. *p < 0.05, **p < 0.01. ***p < 0.001. ns, not significant.

Compound 21 (C21), a DREADD agonist (*66*), was administered 1 hour before photometry recordings to inhibit pIC^→PBN^ neurons. In response to looming stimuli, CGRP^PBN^ activation was significantly reduced in the hM4Di group compared to controls (**Fig. 6, C to F**), with a persistent suppression lasting over 1 min (**Fig. 6 G**). By contrast, CGRP^PBN^ neurons in the hM4Di group responded normally to foot shock (**Fig. 6, H–L**). These findings indicate that pIC input is required for looming-induced CGRP^PBN^ activation but not for foot-shock responses. Given the pIC’s primary role in processing aversive affect, the absence of pIC^→PBN^ input during foot shock may be compensated by alternative nociceptive pathways, such as those involving the spinal cord or PAG.

Further axon tracing of pIC^→PBN^ neurons (**fig. S6**) revealed projections to multiple brain regions, including the CeA, Basomedial amygdalar nucleus (BMA), Parasubthalamic nucleus (PSTN), posterior thalamic regions, and PAG—key areas in emotional regulation and learning (*39*, *67–70*). This suggests that pIC^→PBN^ neurons encode aversive states and broadly relay this information to coordinate behavioral and physiological responses. The observed collateral projections imply that pIC^→PBN^ input contributing to CGRP^PBN^ neuronal responses to looming stimuli may be relayed either directly or indirectly through other regions.

### pIC-to-PBN pathway is necessary for visual threat conditioning but not for foot-shock conditioning

Since pIC^→PBN^ neurons are essential for CGRP^PBN^ activation by visual threats, we tested whether the direct pIC-to-PBN pathway is required for visual threat conditioning. To inhibit this pathway during conditioning, we injected AAV encoding eNpHR3.0-EYFP (or EGFP as a control) into the bilateral pIC and implanted optic fibers above the PBN (**Fig. 7, A and B**). Optogenetic inhibition during visual threat conditioning did not alter immediate freezing but impaired fear memory formation (**Fig. 7, C and D**). While control mice exhibited significantly increased freezing to the tone CS during retrieval, eNpHR3.0-expressing mice showed no significant increase.

**Fig. 7.**
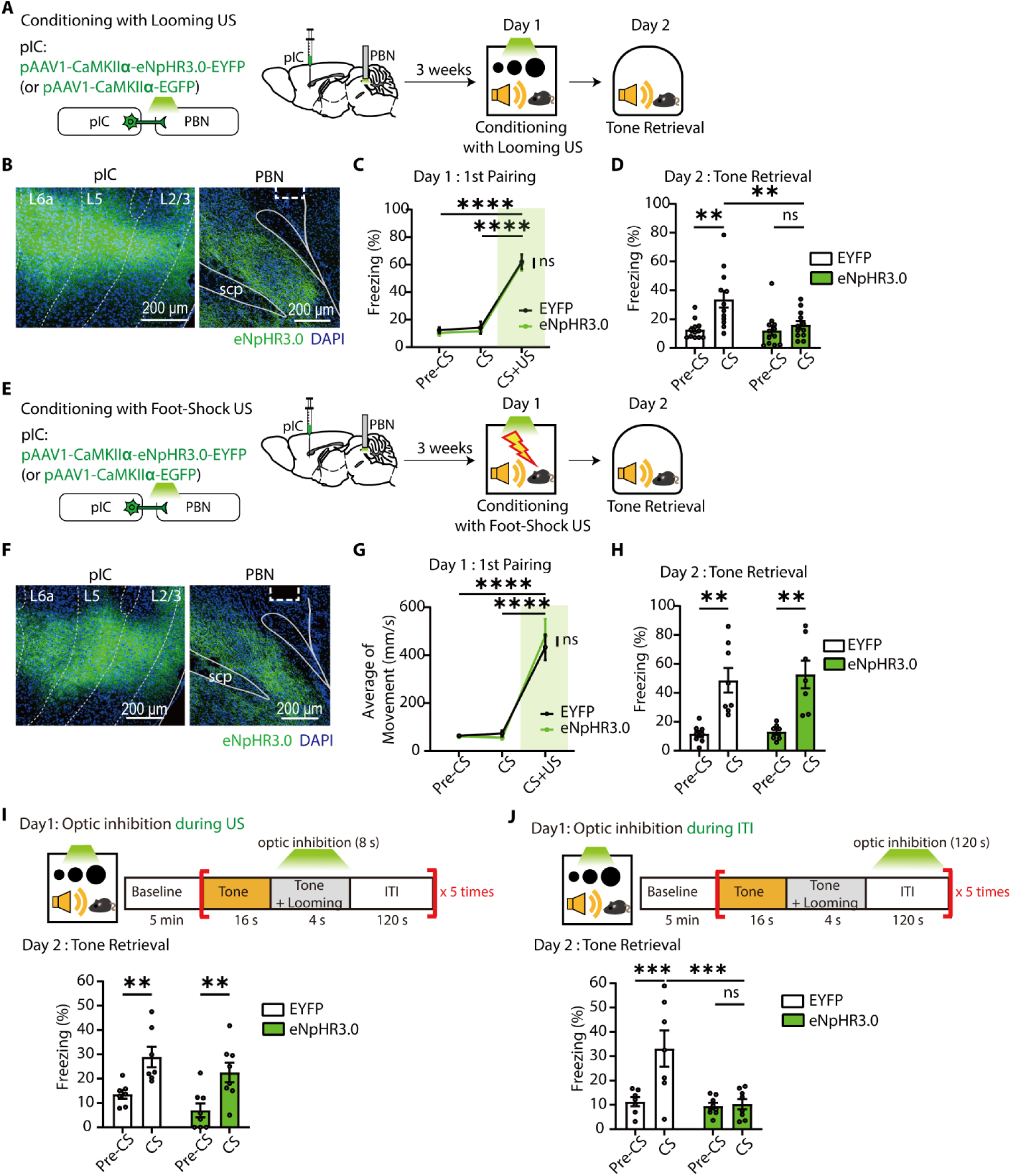
pIC projections to the PBN are required for threat conditioning with looming but not with foot-shock stimuli. (**A**) Schematic of optogenetic inhibition of pIC-PBN circuit during visual threat conditioning. (**B**) Representative images of eNpHR3.0-EYFP expression in pIC neurons (left) and their axon fibers in the PBN (right). The dotted rectangle marks the optic fiber tip. Scale bars, 200 µm. (**C**) Freezing levels during the first CS-US pairing (EYFP, *N*= 12; eNpHR3.0, *N* =12; RM two-way ANOVA, Time, *F_2_*_,44_= 125.6, *P*< 0.0001). (**D**) Optic inhibition of pIC axon terminals in the PBN during conditioning impaired auditory fear memory formation (RM two-way ANOVA, Time × Group, *F*_1,22_= 4.550, *P*= 0.0443). (**E**) Schematic of optogenetic inhibition of pIC-PBN circuit during foot-shock conditioning. (**F**) Representative images of eNpHR3.0 expression in pIC neurons (left) and their axon fibers in the PBN (right). The dotted rectangle marks the optic fiber tip. Scale bars, 200 µm. (**G**) Average of movement during the first pairing of tone CS and foot-shock US on Day 1 (EYFP, *N* = 8; eNpHR3.0, *N* =7; RM two-way ANOVA, Time, *F*_2,26_= 71.34, *P*< 0.0001). (**H**) Optic inhibition of pIC axon terminals in the PBN during foot-shock conditioning did not affect the formation of fear memory (RM two-way ANOVA, Time, *F*_1,13_= 30.91, *P*< 0.0001). (**I**) Optogenetic inhibition of pIC axon terminals in the PBN during US exposure did not impair auditory fear memory formation (EYFP, *N* = 7; eNpHR3.0, *N* =8; RM two-way ANOVA, Time, *F*_1,13_= 38.2, *P*< 0.0001). (**J**) Optogenetic inhibition of pIC axon terminals in the PBN during ITI impaired auditory fear memory formation (EYFP, *N* = 7; eNpHR3.0, *N* =8; RM two-way ANOVA, Time × Group, *F*_1,13_= 12.76, *P*= 0.0034). All data are mean ± s.e.m. **p < 0.01. ***p < 0.001. ****p < 0.0001. ns, not significant.

Next, we examined whether the pIC-PBN pathway is necessary for foot-shock conditioning. For foot-shock conditioning, a 20-s tone CS was presented, with the last 2 s paired with a 0.5-mA foot shock (**Fig. 7, E and F**). This tone-shock pairing was repeated five times while inhibiting pIC-PBN terminals during conditioning. Optogenetic inhibition did not affect defensive escape behavior elicited by the foot shock (**Fig. 7G**). However, unlike visual threat conditioning, inhibition of pIC-PBN pathway had no effect on foot-shock conditioning (**Fig. 7H**). These results indicate that the pIC-PBN pathway serves as a selective US pathway for visual threat conditioning rather than foot-shock conditioning.

Finally, given the sustained responses of both pIC and CGRP^PBN^ neurons to looming stimuli, we investigated the precise timing at which pIC input is required for visual threat conditioning. We selectively inhibited the pIC→PBN pathway either strictly during US exposure or during the ITI (**Fig. 7, I and J**). Inhibition during US exposure had no effect on conditioned freezing (**Fig. 7I**). However, inhibition during the ITI impaired fear memory formation—control mice exhibited increased freezing to the tone CS, whereas eNpHR3.0-expressing mice did not (**Fig. 7J**). These results show that sustained pIC→PBN input is critical for visual threat conditioning, while transient input during US exposure alone is insufficient.

Overall, these findings suggest that pIC neurons provide essential, persistent input to the PBN for visual threat conditioning but are not required for foot-shock conditioning, consistent with our photometry results (**Fig. 6**).

### pIC^→CGRP-PBN^ projections to the PBN induce aversive behavioral states and drive threat conditioning as a US

Our data indicate that the pIC-to-PBN circuit functions as a US pathway for visual threat conditioning. As we hypothesized that this pathway transmits aversive affective information, we examined whether direct activation of the circuit induces aversive affective states and is sufficient to drive threat conditioning. For this purpose, we targeted pIC^→CGRP-PBN^ neurons with channelrhodopsin-2 (ChR2) and optically stimulated their axon terminals in the PBN (**Fig. 8A**). Cre-dependent AAV was injected into the bilateral PBN of *Calca^Cre^* mice, and Flp-dependent AAV encoding ChR2-EYFP (or EYFP alone) was injected into the bilateral pIC. Two weeks later, rabies virus encoding Flp recombinase fused with mCherry was injected into the PBN at the same viral injection sites. Post hoc histological analysis confirmed specific ChR2 expression in pIC neurons, co-expressed with Flp-mCherry (**Fig. 8B**). Starter cells were detected exclusively in the PBN (**fig. S4B**). To assess aversive behaviors, we conducted the real-time place avoidance (RTPA), open-field test (OFT), and elevated plus maze (EPM) tests.

**Fig. 8.**
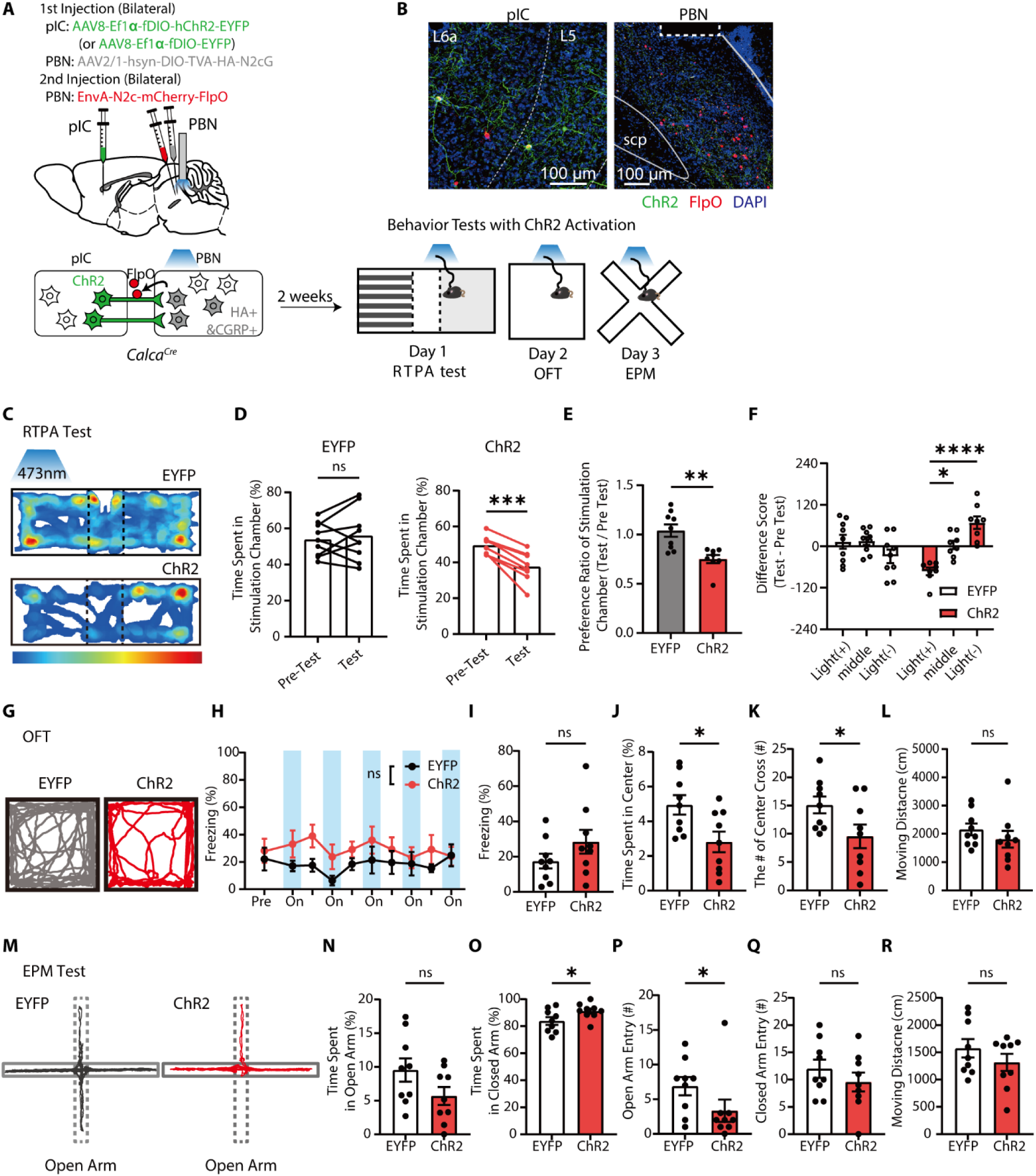
Optogenetic stimulation of pIC^→CGRP-PBN^ projections in the PBN induces anxiogenic behaviors. (**A**) Schematic of viral injections and behavior tests. (**B**) Representative images of ChR2 (green) and/or retrogradely labeled FlpO (red) expression in the pIC (left) and PBN (right). The dotted rectangle marks the optic fiber location. (**C**) Representative heatmaps of time spent in RTPA test chambers during photostimulation. (**D-F**) ChR2 group avoided the photostimulation-paired chamber. (D) Time spent in the photostimulated chamber (Left, EYFP; *N*=9; Two-tailed paired t-test, t_8_= 0.6490, *P*= 0.5345; Right, ChR2; *N*= 8, t_7_= 6.837, *P*= 0.0002). (E) Preference ratio for the photostimulated chamber (Two-tailed Mann Whitney test, U= 6, *P*= 0.0025). (F) Difference score for each chamber (RM two-way ANOVA, Time × Group, *F*_2,30_= 10.77, *P*= 0.0003). (**G**) Representative movement trajectory during the OFT. (**H-L**) ChR2 photostimulation increased anxiety-like behaviors. (H) The photostimulation did not induce a freezing (EYFP, *N*= 9; ChR2, *N*= 9; RM two-way ANOVA, Time × Group, *F*_9,144_= 0.8810, *P*= 0.5439). Averages of freezing levels (I) (Two-tailed unpaired t-test, *t*_16_= 1.379, *P*= 0.1870), time spent in the center zone (J) (*t*_16_= 2.627, *P*= 0.0183), number of center zone crossings (K) (*t*_16_= 2.167, *P*= 0.0456), and moving distance (L) (*t*_16_= 1.131, *P*= 0.2748). (**M**) Representative movement trajectory during EPM. (**N-R**) Time spent in open arms (N) (EYFP, *N*= 9; ChR2, *N*= 9; *t*_16_= 1.406, *P*= 0.0894), time spent in closed arms (O) (*t*_16_= 2.141, *P*= 0.0480), open arm entries (P) (U= 18.5, *P*= 0.0495), closed arm entries (Q) (*t*_16_= 1.012, *P*=0.3267), and moving distance (R) (*t*_16_= 1.129, *P*= 0.2757). Two-tailed Mann Whitney test in (P) and two-tailed unpaired t-test in (N, O, Q, R) were used. All data are mean ± s.e.m. *p < 0.05. **p < 0.01. ***p < 0.001. ****p < 0.0001. ns, not significant.

In the RTPA test, mice were placed in a three-chamber arena where one side was paired with photostimulation of pIC^→CGRP-PBN^ axon terminals in the PBN. The EYFP control group showed no preference between chambers, whereas the ChR2 group avoided the light-paired chamber (**Fig. 8, C to F**). In the OFT, photostimulation was alternated with 60-s intervals (**Fig. 8, G and H**). The ChR2 group did not exhibit increased freezing, suggesting that pIC^→CGRP-PBN^ activation does not induce freezing behavior (**Fig. 8, H and I**). However, ChR2 mice showed increased anxiety, spending less time in the center zone and crossing it fewer times than controls, despite similar total movement distance (**Fig. 8, J to L**). In the EPM test, light stimulation was alternated (2 s on, 2 s off) throughout the trial. ChR2 group spent more time in the closed arms compared to EYFP controls (**Fig. 8O**). Although not statistically significant, they also showed a tendency to spend less time in the open arms (**Fig. 8N**). Additionally, the number of open arm entries was significantly decreased in the ChR2 group (**Fig. 8P**), while total movement distance remained comparable between groups (**Fig. 8R**). These results suggest that activation of pIC^→CGRP-PBN^ projections in the PBN induces aversive affective behavioral states.

We next examined whether photostimulation of pIC^→CGRP-PBN^ projections in the PBN can serve as a US to drive threat conditioning. Using the same viral strategy, we selectively expressed ChR2 in pIC^→CGRP-PBN^ neurons (**Fig. 9, A, B, and fig. S4C**). During conditioning, photostimulation of these projections in the PBN was paired with the tone CS five times. In the tone retrieval test, only the ChR2 group exhibited increased freezing to the tone CS compared to the pre-CS period, while the EYFP control group did not (**Fig. 9C**). These findings demonstrate that pIC^→CGRP-PBN^ neurons transmit aversive affective information to the PBN and that this pathway is sufficient to drive threat conditioning, revealing a distinct circuit for threat learning.

**Fig. 9.**
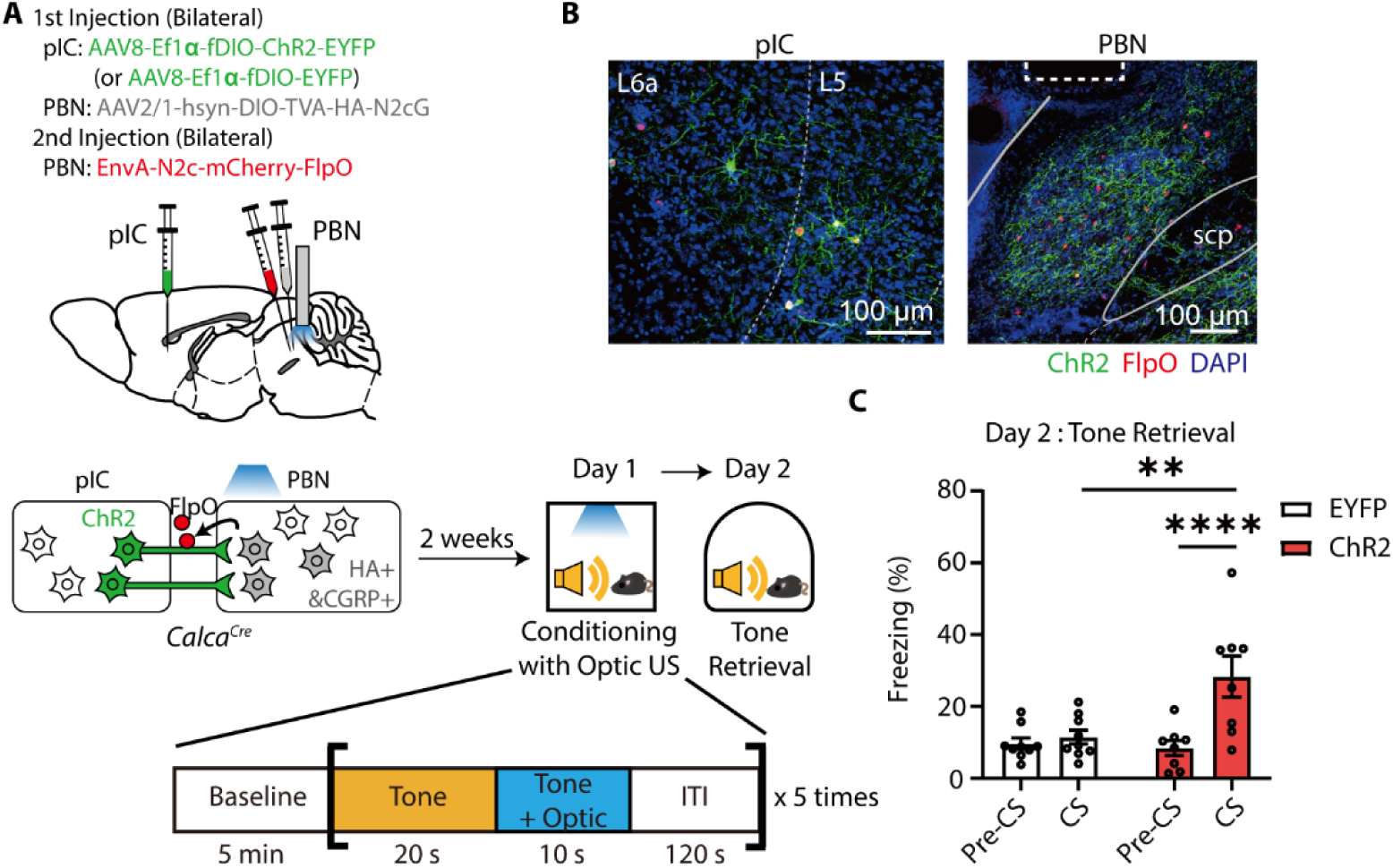
Optogenetic stimulation of pIC^→CGRP-PBN^ projections in the PBN is sufficient to drive formation of fear memory. (**A**) Schematic of a threat conditioning paradigm with optogenetic stimulation of pIC^→CGRP-PBN^ projections in the PBN as s a US. During conditioning, mice were exposed to a 30-s tone, during which the last 10-s tone presentation was overlapped with optic stimulation. This CS-US pairing was repeated 5 times at a 120-s interval. (**B**) Representative images showing the neurons expressing ChR2 (green) and/or retrograde labeled FlpO (Red) in the pIC (left) and ChR2-expressing neurons projections (green) in the PBN (right). Neurons expressing FlpO were also detected in the PBN (right). The dotted rectangle indicates the location of the optic fiber. Scale bar, 100 µm. (**C**) ChR2-expressing mice exhibited increased freezing levels to the conditioned tone during the fear memory retrieval test (EYFP, *N*= 9; ChR2, *N*= 8; RM two-way ANOVA, Time × Group, *F_1_*_,15_= 15.22, *P*= 0.0014). All data are mean ± s.e.m. **p < 0.01. ****p < 0.0001. ns, not significant.

## Discussion

In this study, we established a visual threat conditioning model using looming stimuli as a US in mice. We demonstrated that CGRP^PBN^ neurons are essential for visual threat conditioning and play a broader role in fear learning across both nociceptive and non-nociceptive US. Additionally, we identified a distinct top-down pIC→CGRP^PBN^ circuit as a critical aversive US pathway required for visual threat conditioning. Both pIC^→CGRP-PBN^ and CGRP^PBN^ neurons exhibited threat-specific responses, and the pIC-PBN circuit was necessary for visual threat conditioning but dispensable for foot-shock conditioning. Moreover, activation of pIC^→CGRP-PBN^ projections in the PBN induced aversive affective states and was sufficient to drive fear memory formation. Given the monosynaptic connectivity between pIC and CGRP^PBN^ neurons, these findings suggest that the pIC→CGRP^PBN^ pathway functions as a primary US pathway for non-nociceptive danger signals.

### CGRP^PBN^ neurons as a hub for threat assessment and fear learning

CGRP^PBN^ neurons integrate external and internal signals for comprehensive threat assessment, acting as a general danger-detection hub (*38*, *39*, *51*). They selectively respond to threats like looming and foot shock but not to neutral stimuli (**Fig. 2**, **Fig. 6**), highlighting their role in associative fear learning. After conditioning, their responses to the tone CS increased (**Fig. 2M to O**), suggesting synaptic plasticity enables encoding of learned danger. Both inactivation of CGRP^PBN^ neurons during conditioning and retrieval impaired conditioned freezing (**Fig. 3, G to L**), indicating their necessity for both fear memory formation and recall, reinforcing their central role in visual threat conditioning.

### Distinct pIC-PBN pathway in encoding aversive states

We found that the pIC-PBN pathway transmits aversive affective signals crucial for visual threat conditioning. The pIC^→CGRP-PBN^ neurons selectively respond to both visual and foot-shock threats, but not to neutral stimuli such as flickering (**Fig. 5**), highlighting their role in processing aversive rather than neutral sensory input. Consistent with this, human studies show that the insula is involved in both nociceptive and psychological distress (*58*, *59*). These findings suggest that pIC^→CGRP-PBN^ neurons may constitute a specific population that conveys affective pain signals from multimodal sensory inputs to the PBN. Additionally, optogenetic activation of the pIC^→CGRP-PBN^ pathway induced an aversive state and drove fear memory formation in mice, reinforcing its role in encoding aversive affect (**Fig. 8 and Fig. 9**).

Importantly, inhibition of the pIC-PBN pathway during the ITI impaired fear memory formation (**Fig. 7J**), suggesting that sustained input from the pIC to PBN is required for visual threat conditioning. This prolonged pIC activity may be maintained by interoceptive inputs from physiological changes induced by threats. Indeed, activation of SC parvalbumin-positive (PV+) neurons, which receive direct visual input from retinal ganglion cells (RGCs) and selectively respond to looming threats, increases heart rate (*71*), and CGRP^PBN^ neurons contribute to tachycardia and anxiety-like behaviors (*72*). Given the role of pIC in bottom-up cardiac interoceptive processing (*73*), physiological signals may help maintain long-lasting pIC activity. Alternatively, local recurrent connectivity, involving fast feedback inhibition and slow neuromodulatory transmission (*74*, *75*), or bidirectional interactions with the PBN (*45*, *46*), could contribute to prolonged activation.

In contrast, the pIC-PBN circuit was not essential for foot-shock conditioning (**Fig. 7H**). Additionally, silencing pIC^→PBN^ neurons disrupted CGRP^PBN^ responses to looming but not to foot shock (**Fig. 6**), suggesting that visual threats, which lack sensory pain, primarily engage the pIC for aversive processing. Meanwhile, foot shocks, which involve both affective and sensory pain, can recruit alternative pathways such as the spinal cord and PAG to activate CGRP^PBN^ neurons and support fear learning

### Dissociating innate defensive responses from learned fear

A critical distinction in our study is the separation of innate defensive responses and fear learning processes. CGRP^PBN^ neurons were required for threat conditioning and learned fear responses but not for immediate defensive behaviors to visual threats (**Fig. 3D and fig. S2**). This aligns with studies showing that CGRP+ neurons in the thalamic parvocellular subparafascicular nucleus (CGRP^SPFp^), rather than CGRP^PBN^ neurons, mediate innate freezing to looming threats (*39*). Interestingly, CGRP^PBN^ neurons are required for innate freezing in response to other threat modalities, such as loud tones (*39*), suggesting a modality-specific dissociation in defensive behaviors. Looming stimuli, which mimic fast-approaching objects, likely elicit rapid, reflex-like reactions, whereas fear learning requires a slower risk-assessment process mediated by the PBN.

The SC is a key structure in processing looming threats and coordinating immediate defensive responses. SC PV+ neurons receive direct input from RGCs and specifically respond to looming threats. Additionally, SC neurons transmit looming-relevant inputs to multiple brain regions, including the PBGN, LPTN, PAG, and VTA, which regulate immediate escape and freezing responses (*43*, *44*, *71*, *76*, *77*). Thus, an SC-derived visual pathway, independent of CGRP^PBN^ neurons, is likely responsible for mediating looming-induced innate freezing.

### The presumed role of the SC-PBN pathway in visual threat processing

Although we did not directly assess the function of the SC-PBN pathway, c-Fos analysis revealed increased activity in SC^→CGRP-PBN^ neurons in response to looming stimuli. Since SC neurons detect specific looming patterns in the visual field, the SC-PBN pathway may rapidly transmit potential threat-related visual information to the PBN. It is also possible that SC-PBN pathway is upstream of initial activation of pIC^→CGRP-PBN^ neurons, either directly or via intermediary structures. Given that pIC^→CGRP-PBN^ neurons show increased activity during threat exposure (**Fig. 5**), the pIC likely receives visual threat information rapidly. However, since the retina and SC lack known direct projections to the pIC, visual signals may be relayed from the SC to the pIC via the PBN. Alternatively, the pIC may receive threat-related signals from other intermediary structures such as the thalamus and amygdala.

The functional role of SC-PBN in fear learning remains unknown. Future studies should investigate the SC-PBN pathway’s role in visual threat conditioning and clarify where the neural circuits for innate freezing and threat conditioning diverge.

### Comparison to previous studies and unresolved questions

A prior study reported a failure to achieve Pavlovian fear conditioning with looming stimuli in rats (*78*), which differs from our findings in mice. This discrepancy may stem from differences in experimental conditions, particularly the strong food drive present in the prior study. CGRP^PBN^ neurons integrate both hunger and danger signals, and their suppression by hunger-related inputs (*79*, *80*) may have interfered with aversive learning. Interestingly, the prior study found that combining looming with foot shock successfully impaired food retrieval behavior, suggesting complementary roles for these stimuli in recruiting fear-learning circuits. Future studies should explore how CGRP^PBN^ neurons receive distinct inputs for different types of threats and how their activity is modulated by physiological states.

Future research should examine whether the pIC-PBN circuit serves as a general US circuit for non-nociceptive threats or is specific to visual stimuli. Additionally, identifying the downstream targets of the pIC-PBN circuit responsible for mediating aversive states and associative learning will be crucial. Since the fiber photometry approach lacks single-cell resolution, it remains unclear whether individual pIC^→PBN^ and CGRP^PBN^ neurons respond to both visual and tactile inputs or if distinct subpopulations exist for each modality. Given the role of CGRP^PBN^ neurons in integrating affective signals, investigating the human homologs of this circuit could provide valuable insights into therapeutic targets for anxiety and fear-related disorders.

### Conclusion

Our findings reveal a top-down circuit mechanism by which non-nociceptive visual stimuli induce aversive behavioral states and drive fear memory formation. Identifying this distinct fear memory circuit broadens our understanding of how environmental threats shape fear memory and may inform treatments for threat-related neuropsychiatric disorders, including PTSD.

## Materials and Methods

### Mouse strains

Adult (8-24 weeks) C57BL/6J and heterozygous *Calca^Cre^* (C57BL/6J) mice were used for experiments. Heterozygous *Calca^Cre^* mice were generated by crossing homozygote *Calca^Cre^* mice (JAX #033168) and C57BL/6J wild-type mice. Both male and female mice were used in balance, as we did not observe any noticeable differences in behaviors between male and female. Mice were group housed (3-5 mice per cage) under a 12 h light/dark cycle at a constant humidity (40-60%) and temperature (20-24℃). Food and water were available *ad libitum*. All procedures and protocols were approved by the KAIST Institutional Animal Care and Use Committee (KAIST IACUC, protocol # KA2020-16). All experiments were performed in accordance with the guideline of the KAIST Institutional Animal Care and Use Committee (KAIST IACUC).

### Viruses

AAV1-hSyn-DIO-TeTox-P2A-nls-dTomato (5 × 10^12^ vg/ml), AAV1-hSyn-DIO-tdTomato (8 × 10^11^ vg/ml), AAV1-Ef1α-DIO-eNpHR3.0-EYFP (1 × 10^12^ vg/ml), AAV1-CaMKⅡα-eNpHR3.0-EYFP (2.5 × 10^12^ vg/ml), AAV1-CaMKⅡα-EGFP (2 × 10^12^ vg/ml), AAV8-hSyn-fDIO-mCehrry (8 × 10^11^ vg/ml), AAV8-Ef1α-fDIO-hChR2-EYFP (5 × 10^13^ vg/ml), AAV8-Ef1α-fDIO-EYFP (3 × 10^13^ vg/ml) were packaged.

AAV1-Ef1α-DIO EYFP (2 × 10^13^ vg/ml), AAV8-hSyn-fDIO-hM4Di-mCehrry (2.5 × 10^13^ vg/ml), AAVretro-Ef1α-FlpO (2.3 × 10^13^ vg/ml), AAV1-hsyn-DIO-GCaMP6m (1.9 × 10^13^ vg/ml), AAV8-Ef1α-fDIO-GCaMP6s (1.3 × 10^13^ vg/ml), AAV1-Ef1α-DIO-FlpO (2.1 × 10^13^ vg/ml) were purchased from Addgene. AAVdj-hsyn-fDIO-mGFP-2A-Synaptophysin-mRuby (1.9 × 10^12^ vg/ml) was purchased from Stanford gene vector and virus core (GVVC).

The helper and rabies viruses were obtained from the Viral Vector Core facility of the Kavli Institute for Systems Neuroscience, NTNU. The titers of each virus are as follows: AAV-hsyn-DIO-TVA-HA-N2cG (5 × 10^11^ vg/ml), EnvA pseudotyped SADB19 Rb GFP (1 × 10^11^ vg/ml), EnvA pseudotyped N2c Rb-mCherry-FlpO (4.4 × 10^8^ vg/ml). The helper virus was diluted to 5 × 10^10^ vg/ml in sterile DPBS (Gibco, ThermoFisher) before use for a specific Cre-dependent TVA expression as recommended by a previous paper (*81*).

### Virus packaging

Adeno-associated virus (AAV) was packaged as previously described (*82*). The AAV DNA vector constructs used in this study were either obtained by subcloning of plasmids or purchased from addgene. Briefly, we amplified and purified DNA plasmid by using Maxiprep kit (QIAGEN). The prepared DNA vector was co-transfected with the viral vector (pAd ΔF6 and AAV2/1 or AAV2/8) into HEK293T cells (ATCC, #CRL-3216) by using calcium phosphate precipitation. Seventy-two hours after transfection, cells were harvested, and virus was purified on iodixanol gradient by ultracentrifugation. Viral titers were measured by qPCR (Rotor-Gene Q, QIAGEN).

### Stereotaxic surgeries

Mice were anesthetized with intraperitoneal injection of pentobarbital (83 mg/kg of body weight). Ophthalmic ointment (Liposic gel, BAUSCH and LOMB) was applied to the eyes of the mice to prevent dehydration. Virus was injected with glass capillary filled with water and 2μl of mineral oil (SIGMA, M5904-500ML) at the tip. The injection pipette was placed at the injection site for additional 10 min post-surgery for diffusion of virus.

For manipulation of CGRP^PBN^ neurons, *Calca^Cre^* mice receives bilateral injection of virus (0.45μl per side) using the following coordinate: PBN (AP −5.0mm, ML ±1.4mm, DV −3.55mm). For optogenetic inhibition of CGRP^PBN^ neurons, the optic fibers (Doric lenses, 200μm core diameter, 0.37 NA for optogenetic manipulation) were implanted using the following coordinate: PBN (AP −5.0mm, ML ±1.75mm, DV −3.1mm). For fiber photometry of CGRP^PBN^ neurons, the optic fiber (RWD, 400μm core diameter, 0.5 NA) was implanted unilaterally on the right PBN. The mice underwent behavioral testing after a 2-week recovery period.

For unilateral retrograde tracing, *Calca^Cre^* mice received unilateral injection of helper virus (AAV-hsyn-DIO-TVA-HA-N2cG, 0.45μl) into the right hemisphere of PBN (AP −5.0mm, ML +1.4mm, DV −3.55mm). Two weeks later, rabies virus (EnvA pseudotyped SADB19 Rb GFP, 0.45μl) was injected into the same coordinate. Eight days later, mice were sacrificed for histological analysis. For bilateral retrograde tracing and c-Fos quantification, the injection surgeries were conducted on the bilateral PBN as above. Eight days later, behavioral tests were performed, and mice were sacrificed for histological analysis.

To inhibit terminals of pIC-PBN pathways, C57BL/6J mice received bilateral injection of virus using the following coordinates: pIC (AP −0.4mm, ML ±4.2mm, DV −4.0mm, 0.4μl per side). For optogenetic terminal inhibition, optic fiber (Doric Lenses, 200μm core diameter, 0.37NA) was implanted right above the PBN (AP −5.0mm, ML ±1.75mm, DV −3.1mm). Behavioral experiments were conducted 3 weeks after virus injection.

For fiber photometry of CGRP^PBN^ neurons, AAV1-hsyn-DIO-GCaMP6m was unilaterally injected into the right PBN (AP −5.0mm, ML +1.4mm, DV −3.55mm), and optic fiber (RWD, 400μm core diameter, 0.5 NA) was implanted on the PBN (AP −5.0mm, ML +1.75mm, DV −3.1mm). For fiber photometry of CGRP^PBN^ neurons with hM4Di inhibition of pIC^→PBN^ neurons, we bilaterally injected AAV8-hsyn-fDIO-hM4Di-mCehrry into the pIC and AAVretro-Ef1α-FlpO into the PBN of *Calca^Cre^* mice. Ten days later, AAV1-hsyn-DIO-GCaMP6m was unilaterally injected into the right PBN (AP −5.0mm, ML +1.4mm, DV −3.55mm), and optic fiber (RWD, 400μm core diameter, 0.5 NA) was implanted on the PBN (AP −5.0mm, ML +1.75mm, DV −3.1mm). After 2-week recovery period, the fiber photometry experiments were conducted.

For fiber photometry of pIC^→CGRP-PBN^ neurons, helper virus (AAV-hS-DIO-TVA-HA-N2cG, 0.45μl) was injected into the PBN (AP −5.0mm, ML +1.4mm, DV −3.55mm). Two weeks later, the rabies virus (EnvA pseudotyped N2c Rb-mCherry-FlpO, 0.45μl) was injected into the same site of PBN, and the AAV8-Ef1α-fDIO-GCaMP6s was injected into the ipsilateral pIC (AP −0.4mm, ML +4.2mm, DV −4.0mm, 0.4μl), respectively. Optic fiber (RWD, 400μm core diameter, 0.5 NA) was implanted on the pIC (AP −0.4mm, ML +4.2mm, DV −3.6mm). After a 2-week recovery period, the fiber photometry experiments were conducted.

For ChR2 activation of pIC-CGRP^PBN^ circuit, helper virus (AAV-hS-DIO-TVA-HA-N2cG, 0.45μl) was injected into the bilateral PBN (AP −5.0mm, ML ±1.4mm, DV −3.55mm, 0.45μl), and AAV8-Ef1α-fDIO-hChR2-EYFP was injected into the bilateral pIC (AP −0.4mm, ML ±4.2mm, DV −3.6mm, 0.4μl). Two weeks later, the rabies virus (EnvA pseudotyped N2c Rb-mCherry-FlpO, 0.45μl) was injected into the same site of bilateral PBN, and optic fibers (Doric Lenses, 200μm core diameter, 0.37NA) were implanted on the PBN (AP −5.0mm, ML ±1.75mm, DV −3.1mm). After a 2-week recovery period, the ChR2 activation experiments were conducted.

For axon tracing of PIC^→PBN^ neurons, we injected AAVretro-Ef1α-FlpO into the unilateral PBN, and AAVdj-hsyn-fDIO-mGFP-2A-Synaptophysin-mRuby into the ipsilateral side of pIC. After 3 weeks, the mice were sacrificed for histological analysis.

### Behavioral procedures

#### Looming visual stimuli and behavioral apparatus

The looming visual stimuli were generated using custom software in the LabVIEW (National Instruments) environment and displayed on an overhead LCD monitor (38 cm × 22 cm, 60Hz refresh rate, Intehill). The monitor was positioned 30 cm above the mice and presented a white background. For the looming visual stimuli, a dark disc expanded from 5° (2.6 cm) to 35° (18.9 cm) of visual angle over 300 ms, remained on the screen for 500 ms, and was repeated five times (4 s) during conditioning and ten times (8 s) during the behavioral response test, respectively. For the flickering visual stimuli, a 35° (18.9 cm) dark disc appeared on the monitor for 0.8 s, disappeared for 0.8 s, and this sequence was repeated five times (8 s).

The behavioral arena was surrounded by white acrylic walls, with one of the four walls being transparent (35 cm long, 35 cm wide, 30 cm high). During the experiments, the mouse’s behavior was recorded through the transparent wall using a video camera positioned diagonally over the arena. The video was captured at a frame rate of 30 FPS using LabVIEW software.

#### Innate behavioral responses to looming visual stimuli

All animals were handled for 3 days prior to behavioral tests. To assess innate defensive responses to looming visual stimuli, mice were moved to the behavioral arena from their home cages. After more than 2 min of baseline exploration period, mice were exposed to looming visual stimuli for 8 s (10 repetitions of an expanding dark disc). One minute after the exposure to the looming stimuli, animals were removed from the arena and returned to their home cages. To evaluate the habituation effect to the looming visual stimuli, the immediate response test was conducted for 5 consecutive days. In a single-day habituation test, mice were exposed to the visual stimuli ten times, with a 2-min interval between stimuli. The baseline freezing level was measured for 120 s during the pre-stimulus period before the onset of looming visual stimuli, and the looming-evoked freezing level was measured for 8 s during the visual stimuli presentation period.

#### Open-field test

In the TetTox inhibition experiment, an open field test was performed prior to visual threat conditioning to confirm that there were no differences in basal locomotion and anxiety levels between groups. Mice were placed in an open field arena (45 cm long, 45 cm wide, 30 cm high) and allowed to freely explore for 15 min. Freezing levels, total distance traveled, and time spent in the center area of the arena were measured.

#### Visual threat conditioning and fear memory retrieval test

All animals were handled for 3 days prior to behavioral tests. The conditioning chamber was a white acrylic box with one transparent wall, measuring 35 cm in length, 35 cm in width, and 30 cm in height. The floor was covered with paper pads. After 5 min of baseline exploration, mice were exposed to a 20-s auditory tone (CS, 2.7 kHz, 80 dB) that was co-terminated with 4-s looming visual stimuli (US, a dark disc expanding five times). After five CS-US pairings with a 2-min interval, mice were removed from the conditioning chamber. In the unpaired conditioning protocol, the same auditory tone was presented but with random intervals, ranging from 15 to 30 s after the looming visual stimuli. The baseline freezing level was measured for 120 s during the pre-CS period immediately before the tone onset. Freezing levels for the CS and CS+US periods were measured during the first 16 s and last 4 s of CS-US pairing, respectively.

One day after conditioning, an auditory fear memory test was conducted. The mice were placed in a test chamber with a shifted context: a semicircular arena, striped walls, and a white acrylic floor (35 cm long, 35 cm wide, 30 cm high). After more than 2 min of exploration, the conditioned tone was presented for 20 s. Freezing behavior was measured during the pre-CS period (120 s) and the CS period (20 s).

Contextual fear memory test was conducted the following day. Mice were re-exposed to the conditioning chamber for 5 min, and freezing behavior was measured during the first 2 min to assess contextual fear memory.

#### Foot-shock threat conditioning and fear memory retrieval test

All mice underwent 3 days of handling prior to behavioral testing. For foot-shock conditioning, mice explored a conditioning chamber equipped with a metal grid floor (Med Associates, 25 cm long, 30 cm wide, 20 cm high) for 5 min. Subsequently, they were exposed to a 20-s auditory tone (CS, 2.7 kHz, 80 dB) that was co-terminated with a 2-s foot-shock stimulus (US, 0.5 mA). The CS-US pairing was repeated 5 times with a 2-min interval. The average of baseline movement was measured for 120 s during the pre-CS period immediately before the onset of foot-shock. Movement for the CS and CS+US periods was measured for the first 16 s and the last 4 s during CS-US pairing, respectively.

One day after foot-shock conditioning, an auditory fear memory test was conducted. Mice were placed in a test chamber with a shifted context: a semicircular arena, striped walls, and a white acrylic floor (Med Associates, 25 cm long, 30 cm wide, 20 cm high). After more than 2 min of exploration, the conditioned tone was presented for 20 s. Freezing behavior was measured during the pre-CS period (120 s) and the CS period (20 s).

#### Optogenetic inhibition behavior experiments

All mice underwent three days of handling before behavioral testing. To habituate them to the optical manipulation procedure, they were tethered to fiber-optic patch cords for five minutes per day over three consecutive days.

For optogenetic eNpHR3.0 inhibition during conditioning, optical inhibition (561 nm, continuous; 3–5 mW at the fiber tip for soma inhibition, 10 mW for terminal inhibition) began two seconds before the first US exposure and continued until the end of the experiment.

For optogenetic inhibition during the tone retrieval test, optical inhibition was applied throughout the 20-second presentation of the tone-CS and terminated immediately afterward.

For optogenetic inhibition during the US exposure experiment, inhibition was initiated two seconds before US onset and terminated two seconds after US offset, lasting a total of eight seconds per trial.

For optogenetic inhibition during the ITI, inhibition started immediately after US exposure and ended just before the next US exposure, lasting two minutes per trial.

#### Optogenetic stimulation behavior experiments

All mice underwent 3 days of handling before behavioral testing. Additionally, mice were tethered to fiber-optic patch cords for 5 min per day over 3 consecutive days to habituate them to the optic manipulation procedure.

To assess the anxiogenic effects of stimulating the pIC-CGRP^PBN^ pathway, we conducted the RTPA test, open-field test, and elevated plus maze (EPM) test sequentially. In the RTPA test, the three-chamber arena was used (25 cm long, 75 cm wide, and 25 cm high; two side chambers with 30 cm wide each, and a middle chamber with 15 cm wide). One of the side chambers had a striped wall and metal grid floor, while the other had a checkered wall and punched metal floor. During the test, mice explored the arena for total 30 min.

The first 10-min was used to determine a baseline chamber preference between the two side chambers. The chamber with more preference was selected for pairing with photostimulation. During the last 10 min, optical stimulation (473 nm, 20 Hz, 10 ms pulse, 10 mW at the fiber tip) was applied only when mice entered the selected side chamber. To assess the avoidance behavior induced by photostimulation, the time spent in each side chamber during the pre-test (the first 10 min) and test period (the last 10 min) was compared. The preference ratio was calculated by dividing the time spent in the photostimulation-paired chamber during the test by the time spent in the same chamber during the pre-test. The difference score was calculated by subtracting time spent in each chamber (left, middle, and right) during the pre-test from during the test. One mouse that spent more than 70% of the time in one of the two side chambers during the pre-test period was excluded from the analysis.

In the open-field test, to evaluate the effect of optical stimulation on immediate freezing and anxiogenic behaviors, mice were allowed to explore an open-field arena (45 cm x 45 cm x 30 cm). After 2-min exploration, the mice were exposed to a 30 s-photostimulation (473 nm, 20 Hz, 10 ms pulse, 10 mW at the fiber tip) session five times with a 60-s interval. Measures included freezing behavior, time spent in the center zone, the number of center crossings, and total distance traveled. The center zone was defined as a 20 cm square area in the center of the arena.

In the EPM test, the arena was a cross-shaped acrylic structure with 65 cm in length and 5 cm in width. Two closed arms were enclosed by 16 cm high white acrylic walls, while the open arms have 0.5 cm high transparent acrylic walls. Mice were allowed to explore the EPM arena freely for 10 min, during which they were exposed to photostimulation with a 2-s on and 2-s off cycle. To evaluate anxiety behavior, the time spent in the open and closed arms, the number of entries into each arm, and the total distance traveled during the 10-min test were measured.

For optic conditioning, mice were allowed to explore a conditioning chamber equipped with a metal grid floor (Med Associates, 25 cm long, 30 cm wide, 20 cm high) for 5 min. Subsequently, they were exposed to a 30-s auditory tone (CS, 2.7 kHz, 80 dB) that was co-terminated with a 10-s photostimulation (US, 473 nm, 20 Hz, 10 ms pulse, 10 mW at the fiber tip). The CS-US pairing was repeated 5 times with a 2-min interval. One day after conditioning, mice were tested for auditory fear memory. Mice were placed in a test chamber with a shifted context: a semicircular arena, striped walls, and a white acrylic (Med Associates, 25 cm long, 30 cm wide, 20 cm high). After more than 2 min of exploration, the conditioned tone was presented for 30 s. Freezing behavior was measured during the pre-CS period (120 s) and the CS period (30 s).

#### c-Fos imaging behavior experiments

To assess c-Fos levels in the projection neurons upstream to CGRP^PBN^ neurons in response to looming visual stimuli, behavioral experiments were conducted 8 days after rabies virus injection. Mice were allowed to freely explore the test arena for 5 min before being exposed to visual stimuli (either looming visual stimuli for 8 s with 10 expansions or flickering visual stimuli for 8 s with 5 flickers) twice at a 120-s interval. The mice were then returned to their home cages and sacrificed 90 min after the first stimulus exposure.

### Fiber photometry

For fiber photometry, a 400 µm diameter, 0.48 NA patch cord was used to collect fluorescence emission. A real-time signal processor [Tucker-Davis Technologies (TDT), RZ5P) controlled the 470 nm (Thor Labs, M470F3) and 405 nm (Doric, LEDC1-405_FC) LEDs and simultaneously recorded GCaMP fluorescence emission. The 470 nm LED was sinusoidally modulated at 331 Hz to excite GCaMP, while the 405 nm LED was modulated at 531 Hz to monitor the isosbestic signal. The 470nm and 405nm light passed through a bandpass filter in a mini cube (Doric, FMC5), and into the optic fiber connected to the brain of mice. The emitted GCaMP6m signal was detected by a photoreceiver (New Focus, Model 2151) and digitized at 1017.3 Hz by the RZ5P processor. Fluorescence measurements (<6 Hz) were extracted from Synapse software (TDT), and further data processing was performed in MATLAB. ΔF/F was calculated by applying a least-squares linear fit to the 405 nm signal, aligning it with the 470 nm signal (fitted 405nm), and then computing ΔF/F as (470 nm -fitted 405 nm)/fitted 405 nm.

Behavioral experiments were recorded at 30 FPS, with stimulus delivery and video capture controlled by a custom LabView program. To synchronize photometry recording with behavioral events (e.g., tone, visual stimuli, shock), digital signals generated by LabView were recorded by the RZ5P processor.

To record calcium activity of CGRP^PBN^ neurons in response to flickering and looming visual stimuli, mice were placed in the looming test chamber and allowed to explore freely for 5 minutes before stimulus presentation. First, they were exposed to flickering stimuli, followed by looming stimuli after a 5-minute interval.

To examine calcium activity of CGRP^PBN^ neurons while chemogenetically inhibiting pIC^→PBN^ neurons, DREADD agonist 21 dihydrochloride (C21; Tocris, #6422; 0.2 mg/ml working solution, 10 mg/ml stock) was administered via intraperitoneal injection at a dose of 1 mg/kg. One hour after C21 injection, mice were placed in the looming test chamber, where they explored for 5 minutes before being exposed to two looming stimuli (8 s) at a 120-second interval. After a 10-minute rest, mice were transferred to a foot-shock chamber, where they explored for 5 minutes before receiving two foot-shocks (2 s, 0.5 mA) at a 120-second interval. To compare ΔF/F across animals during stimulus exposure, ΔF/F was normalized to a Z-score, calculated as [(ΔF/F – Mean of ΔF/F_baseline_)/Std of ΔF/F_baseline_], where the baseline mean and standard deviation were computed from –5 to 0 s before stimulus onset.

To record the calcium response of pIC^→CGRP-PBN^ neurons to visual or foot-shock stimuli, mice were first placed in the looming test chamber, where they explored freely for five minutes before being exposed to looming visual stimuli (or flickering stimuli). The looming (or flickering) stimuli presented twice at a 120-s interval. After a 10-minute rest period, mice were transferred to the foot-shock chamber, where they explored freely for five minutes before receiving two foot-shocks (2 s, 0.5 mA) at a 120-s interval. As pIC^→CGRP-PBN^ neurons exhibited sustained calcium activity in response to both looming and shock stimuli, only the first 60 seconds of recording data from stimulus onset were included in the analysis. To minimize the effect of natural fluctuations in pIC neuronal activity, the baseline for Z-scoring was computed from the 60-second period preceding stimulus onset (–60 to 0 s). To determine the peak and the time to half of the peak in calcium activity, Z-scored calcium activity was filtered using a 1 Hz low-pass filter. The maximum peak was identified within the 0–100 s time window following stimulus onset. After determining the peak time, the time point at which calcium activity declined to half of the peak value was identified. Since pIC^→CGRP-PBN^ neurons exhibited persistent but distinct response dynamics to looming and shock stimuli, we analyzed their responses based on the half-peak time window: 0–60 s for looming and 0–20 s for shock

### Analysis of behaviors

Freezing was defined as a state in which the mouse remained completely immobile except for respiratory movements. Running was defined as when the centroid of the mouse body moved at a velocity of 40 cm/s or greater. All other behaviors were classified as no response. The centroid of the subjects was analyzed to determine position, calculate velocity, and track movement trajectories. Centroid was detected by using custom MATLAB scripts. Brightness thresholds were manually set to differentiate the pixels corresponding to the mouse. Perspective distortion from the camera lens was corrected using the *tform* function in MATLAB. Velocity was calculated from the centroid data and smoothed using a mean filter.

### Histology

For histological verification of virus expression and optic fiber placement, brain sections were prepared from the sacrificed mice after the end of behavioral experiments. Mice were anesthetized with 2.5% avertin by intraperitoneal injection, and they were perfused transcardially with phosphate-buffered saline (PBS) and then fixed with ice cold 4% paraformaldehyde (PFA). After perfusion, brain samples were stored in 4% PFA overnight for post-fixation and sliced in a 40μm thickness afterward by using a vibratome (VT-1200S, Leica Microsystems).

For a retrograde tracing and c-Fos staining, mice underwent behavioral experiments 8 days after the rabies virus injection and were sacrificed 90 min after exposure to visual stimuli. Mice were anesthetized with pentobarbital (83 mg/kg of body weight) by intraperitoneal injection, perfused transcardially with phosphate-buffered saline (PBS), and then fixed with ice cold 4% paraformaldehyde (PFA). After perfusion, brain samples were stored in 4% PFA overnight for post-fixation. Fixed brain samples were dehydrated in a series of 10%, 20%, and 30% filtered sucrose solutions (#S0389, Sigma-Aldrich) until they sank to the bottom of vials. Dehydrated brain samples were attached on a specimen disc with OCT compounds (#4583, Scigen) at −20 ℃. Brains were coronally sliced in a 40μm thickness by using a Cryostat (Leica CM1850, Leica Biosystems).

Brain sections were mounted on a gelatin-coated slide glass with a Vectashield antifade mounting media (h-2000, Vector Laboratories). Images were taken for histological verification by using a confocal microscope (LSM880, Carl ZEISS) or slide scanner (ZEISS Axio Scan.Z1, Carl ZEISS). By reference to the Mouse Brain Atlas, the mice who showed off-target viral expression in surrounding areas or displacement of ferrules were excluded.

### Immunohistochemistry

For immunostaining, brain sections were washed three times with PBS and incubated in blocking solution (2% goat serum, 0.1% BSA, and 0.2% Triton X-100 in PBS) for 1 hour on an orbital shaker (126 rpm). Brain sections were incubated in a primary antibody solution on the orbital shaker (126 rpm) overnight at room temperature (RT). After brain sections were washed four times with PBS, sections were then incubated in a secondary antibody solution on the shaker (126 rpm) for 2 hours. Subsequently, the brain sections underwent four additional washes with PBS. Immunostained brain sections were transferred to gelatin-coated slides using a brush. Sections were stained with DAPI (0.5μg/ml in PBS; #D9542, Sigma-Aldrich) for 20min at RT. The slides were rapidly immersed in a distilled water and allowed to air-dry completely in a light-protected environment. Coverslip was mounted with a Vectashield antifade mounting media (h-2000, Vector Laboratories).

The dTomato fluorescence signals in neurons injected with AAV-TetTox-P2A-nls-dTomato virus were amplified with mouse anti-DsRed primary antibody (1:500; #SC-390909, Santacruz) and Alexa Fluor 594-conjugated goat anti-mouse secondary antibody (1:2000; #A-11005, Molecular Probes). The EYFP fluorescence signals in neurons injected with AAV-CaMKⅡα-eNpHR3.0-EYFP virus were amplified with rabbit anti-GFP primary antibody (1:5000; #ab290, abcam) and Alexa Fluor 488-conjugated goat anti-rabbit secondary antibody (1:2000; #A-11008, Molecular Probes). The antibodies used for HA staining were rabbit anti-HA (1:1000; #3724, Cell signaling) and Alexa Fluor 647-conjugated goat anti-rabbit (1:2000; #A-21244, Molecular Probes). The antibodies used for c-Fos staining were rabbit anti-cFos (rabbit anti-cFos; 1:2000; #226008, Synaptic systems) and Alexa Fluor 594-conjugated goat anti-rabbit (Alexa Fluor 594-conjugated goat anti-rabbit; 1:2000; #A-11037, Molecular Probes). In axon tracing experiment, mGFP and synaptophysin-mRuby were amplified with rabbit anti-GFP primary antibody (1:5000; #ab290, abcam), mouse anti-RFP primary antibody (1:2000; # 200-301-379; Rockland), Alexa Fluor 488-conjugated goat anti-rabbit secondary antibody (1:2000; #A-11008, Molecular Probes), and Alexa Fluor 594-conjugated goat anti-mouse (1:2000; #A-11005, Molecular Probes).

For anterograde tracing of the pIC^→PBN^ neurons, Sudan black B (SBB) staining was performed. After DAPI staining, the samples were stained with 0.1% SBB for 20 minutes at room temperature. The sections were briefly rinsed with 70% ethanol for 30 seconds, followed by a 5-minute rinse in PBS. The 0.3% SBB stock solution (in 70% ethanol) was stirred overnight in the dark and filtered using a 0.22μm filter. The 0.1% SBB working solution was freshly prepared before use.

### Cell counting analysis

To map the location of retrogradely labeled GFP+ cells throughout the whole brain, coronal brain tissues were collected along AP axis (from anterior part AP +1.8 mm to posterior part AP: −5.5 mm) in a 120-µm distance. For the brain-wide tracing, images of all brain sections were captured using a Slide scanner (ZEISS Axio Scan.Z1, Carl ZEISS). The number of DAPI+ and GFP+ cells in the brain sections were then counted by using the AMaSiNe automatic cell counting software (*83*) by registering brain images to the Allen Brain Atlas Common Coordinate Framework (CCFv3) (*84*).

To assess the overlap between c-Fos+ and GFP+ cells, we conducted c-Fos immunostaining, and optical sections (2 μm per section) were obtained using confocal microscopy with Z-stack analysis. DAPI+, GFP+, and c-Fos+ cells were detected using the AMaSiNe cell counting algorithm (*83*). Regions of interest (ROI) were then defined in each target area, and the overlap between c-Fos+ and GFP+ cells was manually counted in a blinded manner. The AP ranges for c-Fos quantification in each target brain area are: pIC (AP −0.2 to −0.6 mm), CeA (AP −1.2 to −1.6 mm), PAG (AP −3.5 to −3.9 mm), SC (AP −3.5 to −3.9 mm).

To compare CGRP^PBN^-projecting neurons in the aIC and pIC, we counted GFP+ cells in the aIC (AP: +1.0 to 0 mm) and pIC (AP: 0 to −1.0 mm).

### Statistical analysis

GraphPad Prism 10.0.2 was used for graph generation and statistical analyses. The Shapiro-Wilk test was used to assess data normality. For datasets with a small sample size (N < 7), the Kolmogorov-Smirnov test was conducted, and data distribution was additionally evaluated using QQ-plots. Paired or unpaired Student’s t-tests were performed to compare normally distributed datasets of experimental and control groups depending on experimental design. Wilcoxon matched-pairs signed rank test or Mann Whitney test were performed to compare non-normally distributed datasets of experimental and control groups. One-way ANOVA and repeated-measures two-way ANOVA, followed by Sidak’s post-hoc test, were used to analyze behavioral tasks depending on the experimental design. Differences were considered statistically significant if the p-value was below 0.05. Error bars represent the s.e.m.

## Acknowledgments

We thank all members of the Memory Biology laboratory for helpful comments, discussions, and constructive suggestions. We thank the KAIST Bio-Core Center and the KAIST Analysis Center for Research Advancement (KARA) for use of confocal microscopes and assistance in performing imaging experiments.

## Author contributions

Conceptualization: J.H., J-H.H.

Data curation: J.H., J-H.H.

Formal analysis: J.H., B.S., J-H.H.

Funding acquisition: J-H.H.

Investigation: J.H., B.S.

Methodology: J.H., J-H.H.

Project administration: J-H.H.

Resources: J-H.H.

Software: J.H.

Supervision: J-H.H.

Validation: J.H., B.S., J-H.H.

Visualization: J.H., J-H.H

Writing—original draft: J.H., J-H.H.

Writing—review & editing: J.H., B.S., J-H.H.

## Data and materials availability

All data needed to evaluate the conclusions in the paper are present in the paper and/or the Supplementary Materials.

---

**fig. S1.**
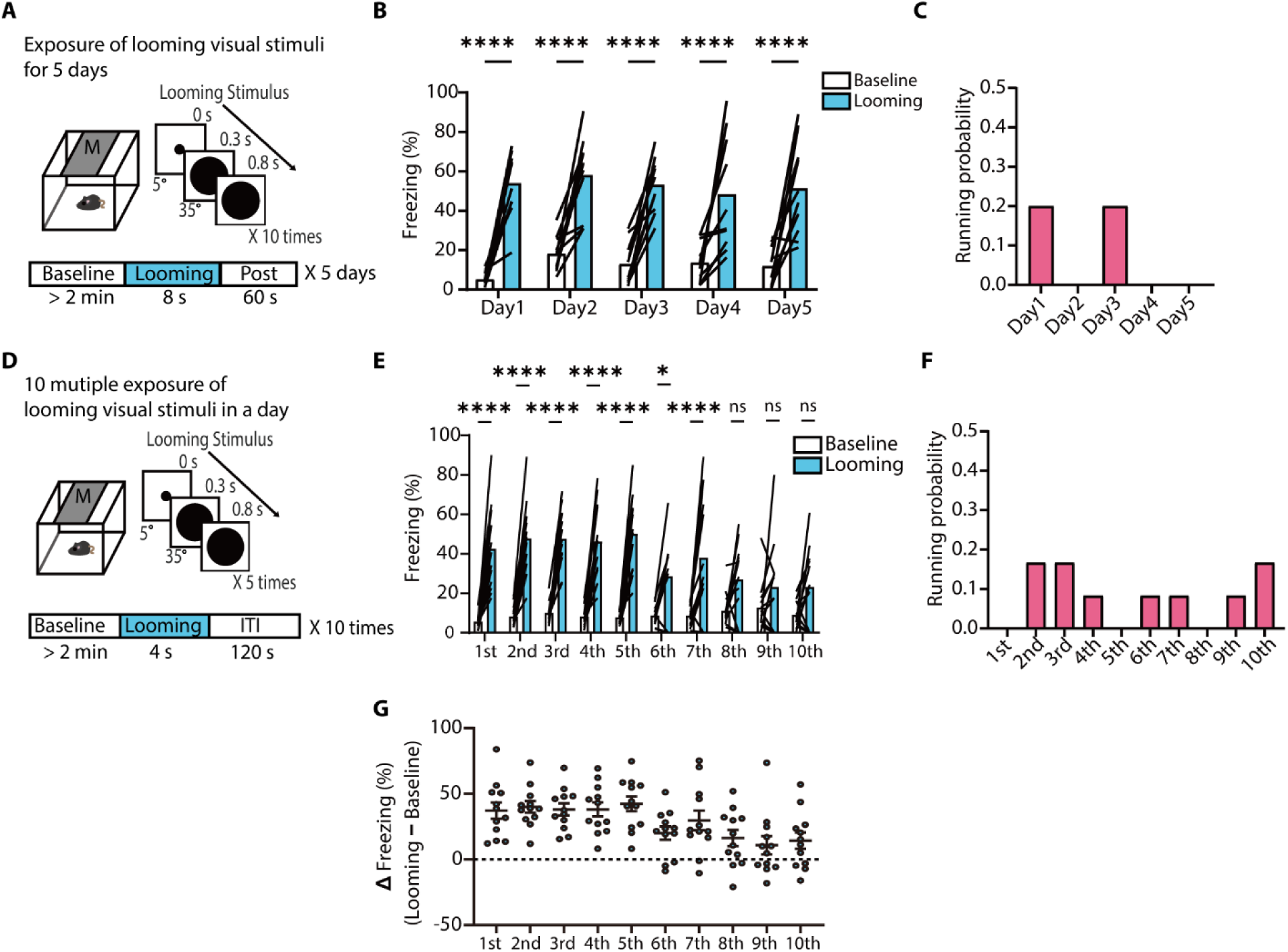
CGRP^PBN^ neuronal activity in response to flickering and looming visual stimuli (related to Figure 1). **(A)** Schematic of repeated behavioral tests for innate defensive responses to looming visual stimuli over 5 days. Each day, mice were exposed to 10 consecutive looming stimuli (8 s per trial). **(B)** No significant habituation to daily looming stimuli over 5 days. Freezing responses to looming stimuli remained consistent (*N* = 10 mice; RM two-way ANOVA, Stimulus, *F*_1,18_= 72.07, *P*< 0.0001). **(C)** Probability of running responses to daily looming stimuli over 5 days, calculated as the number of mice exhibiting running behavior divided by the total number of mice. **(D)** Schematic of a behavioral test for habituation of the innate freezing response to looming stimuli. A set of 5 looming stimuli (4 s each) was repeated 10 times with a 120-s intertrial interval. **(E)** Habituation of innate freezing response. Freezing was measured during each 4-s looming stimuli (*N* = 12 mice; RM two-way ANOVA, Trials × Stimulus, *F*_9,198_= 4.198, *P*< 0.0001). **(F)** Probability of running event to the 10-repeated looming stimuli. **(G)** Difference in freezing levels between baseline and looming stimuli (RM one-way ANOVA, *F*_9,99_= 4.598, *P*< 0.0001). All data are presented as mean ± s.e.m. *p < 0.05, ****p < 0.0001. ns, not significant.

**Fig. S2.**
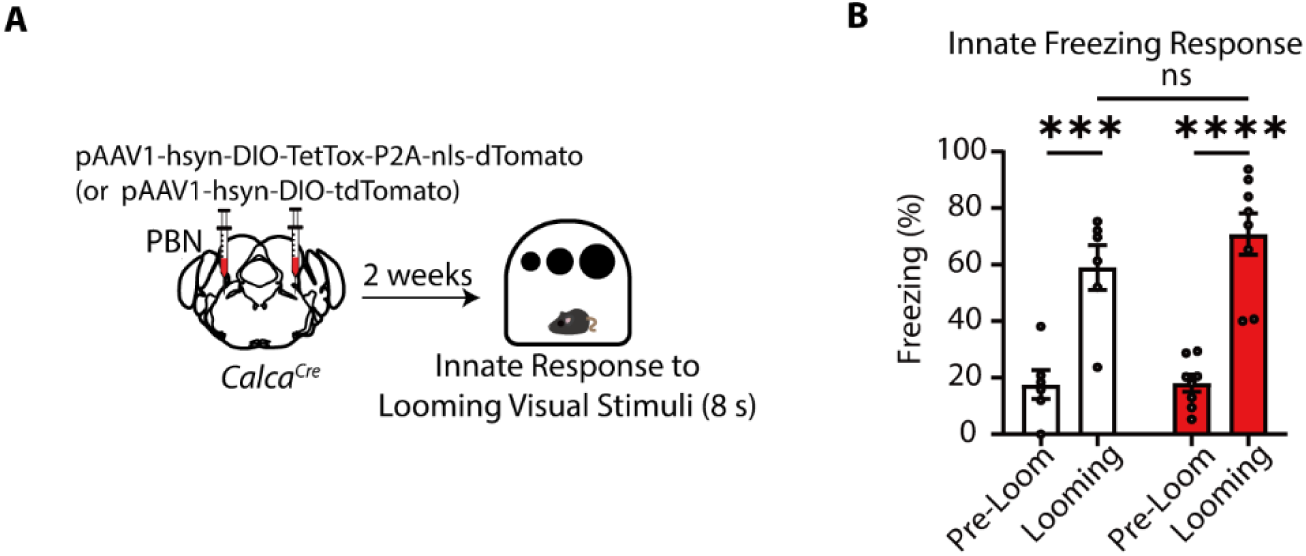
TetTox inhibition of CGRP^PBN^ neurons did not impair innate defensive behavioral responses to looming visual stimuli (related to Figure 3). **(A)** Schematic of innate freezing response test with TetTox-mediated inhibition of CGRP^PBN^ neurons. **(B)** TetTox inhibition of the CGRP^PBN^ neurons did not affect innate freezing response to looming visual stimuli (tdTomato group, *N*= 6 mice; TetTox group, *N*= 8 mice; RM two-way ANOVA, Time, *F*_1,12_= 68.01, *P*< 0.0001). All data are mean ± s.e.m. ***p < 0.001, ****p < 0.0001. ns, not significant.

**Fig. S3.**
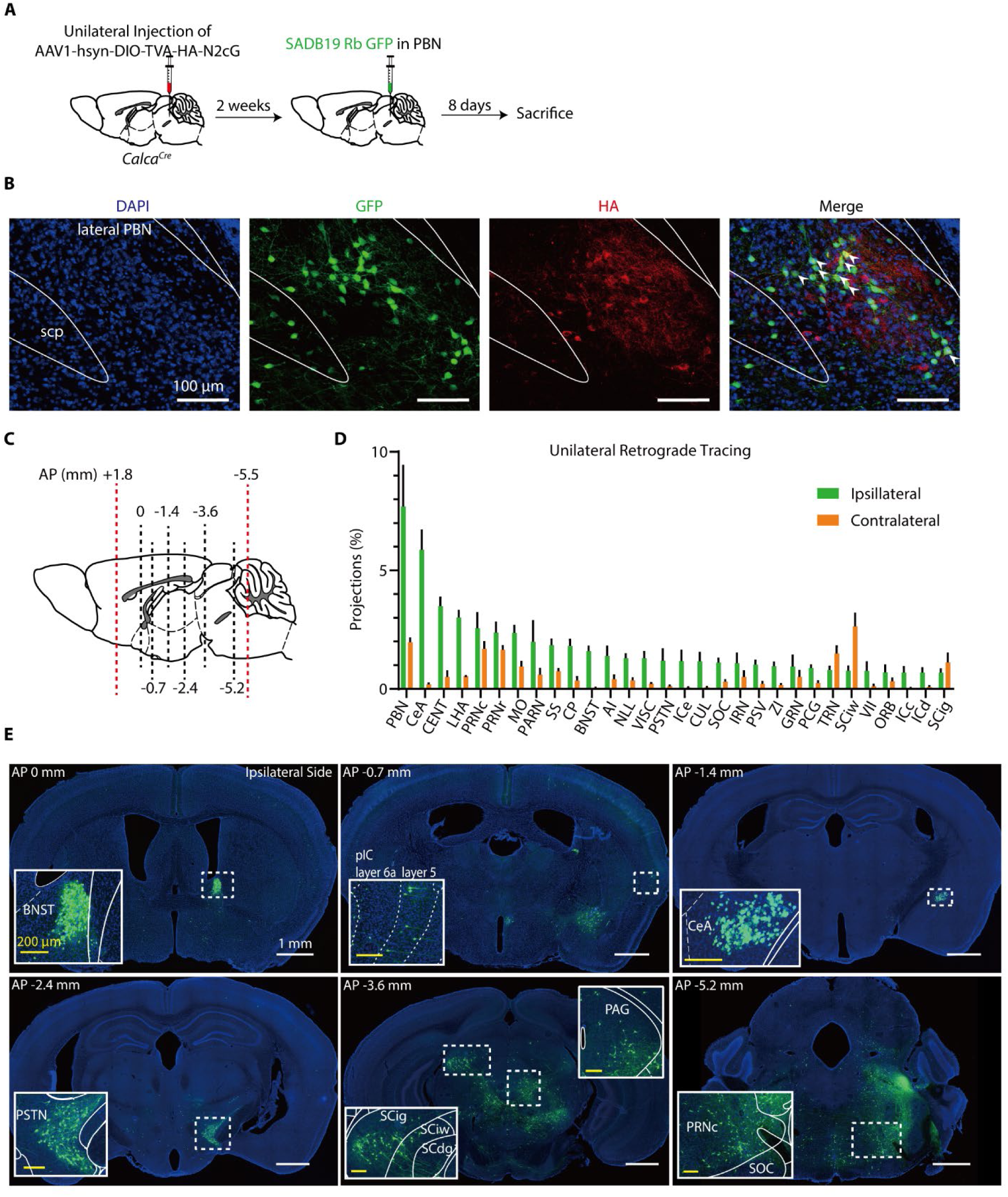
Brain-wide monosynaptic retrograde tracing of CGPR^PBN^ neurons (related to Figure 4). **(A)** Schematic of unilateral monosynaptic retrograde tracing of CGRP^PBN^ neurons. Unilateral injections of AAV2/1-hsyn-DIO-TVA-HA-N2cG into the PBN of *Calca^Cre^* mice were followed by unilateral injections of SADB19-GFP rabies virus into the same site 2 weeks later. **(B)** Representative confocal microscopic images of coronal sections showing neurons expressing GFP and/or haemagglutinin (HA) tag in the PBN. White arrows indicate double-labeled starter cells (GFP+ & HA+). Scale bar, 100 µm. **(C)** AP range of coronal sections collected for imaging analysis (AP, +1.8 mm to −5.5 mm, indicated by dotted red lines) and AP coordinates of representative images shown in (E) (dotted black lines). **(D)** Distribution of retrogradely labeled neurons across the top 30 brain regions with the highest number of projections. **(E)** Representative images of coronal sections showing GFP-labeled retrogradely traced neurons in each analyzed target region. Insets show magnified views of the dotted boxes. Scale bars: 1 mm (white bar) and 200 µm (yellow bar in insets). **Abbreviations**: Agranular insular area (AI); Bed nuclei of the stria terminalis (BNST); Caudoputamen (CP); Central amygdalar nucleus (CeA); Central lobule (CENT); Culmen (CUL); Facial motor nucleus (VII); Gigantocellular reticular nucleus (GRN); Inferior colliculus, central nucleus (ICc); Inferior colliculus, dorsal nucleus (ICd); Inferior colliculus, external nucleus (ICe); Intermediate reticular nucleus (IRN); Lateral hypothalamic area (LHA); Nucleus of the lateral lemniscus (NLL); Orbital area (ORB); Parabrachial nucleus (PBN); Parvicellular reticular nucleus (PARN); Parasubthalamic nucleus (PSTN); Pontine central gray (PCG); Pontine reticular nucleus, caudal part (PRNc); Pontine reticular nucleus (PRNr); Principal sensory nucleus of the trigeminal (PSV); Somatomotor areas (MO); Somatosensory areas (SS); Superior colliculus, motor related, intermediate gray layer (Scig); Superior colliculus, motor related, intermediate white layer (Sciw); Superior olivary complex (SOC); Tegmental reticular nucleus (TRN); Visceral area (VIS); Zona incerta (ZI).

**Fig. S4.**
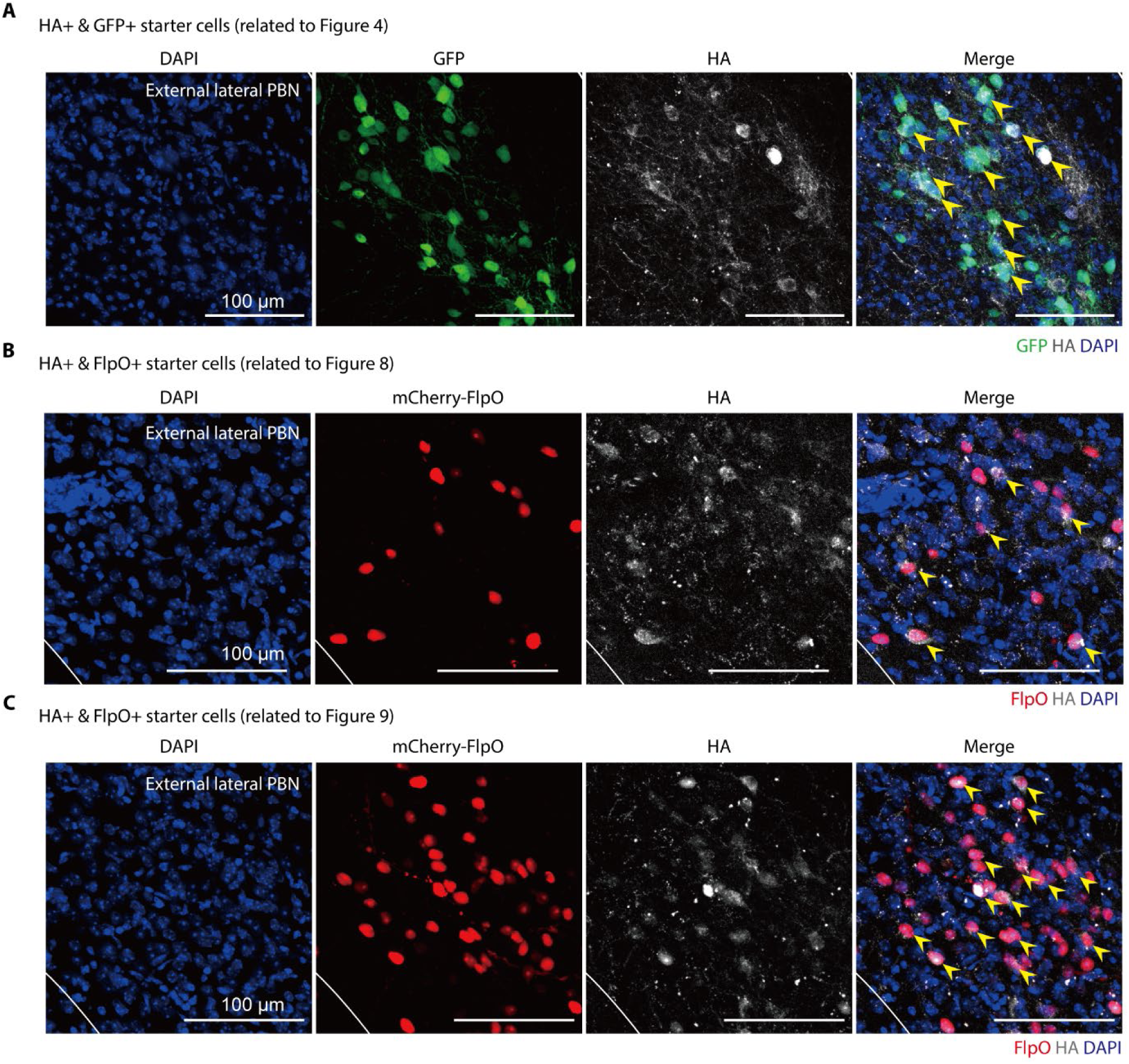
Images of starter cells in the PBN in the monosynaptic retrograde labelling experiments (related to Figure 4, 8, and 9). **(A)** Representative confocal microscopic images of coronal sections showing the neurons double-labeled for GFP and HA tag in the PBN. Yellow arrow head indicates double-labeled starter cells (GFP+ & HA+) in Figure 4. (**B, C**) Representative confocal microscopic images of coronal sections showing the neurons expressing FlpO fused with mCherry and HA tag in the PBN, related to the experiments in Figure 8 (B) and Figure 9 (C). Yellow arrow head indicates double positive starter cells (mCherry-FlpO+ & HA+). Scale bars, 100 µm.

**Fig. S5.**
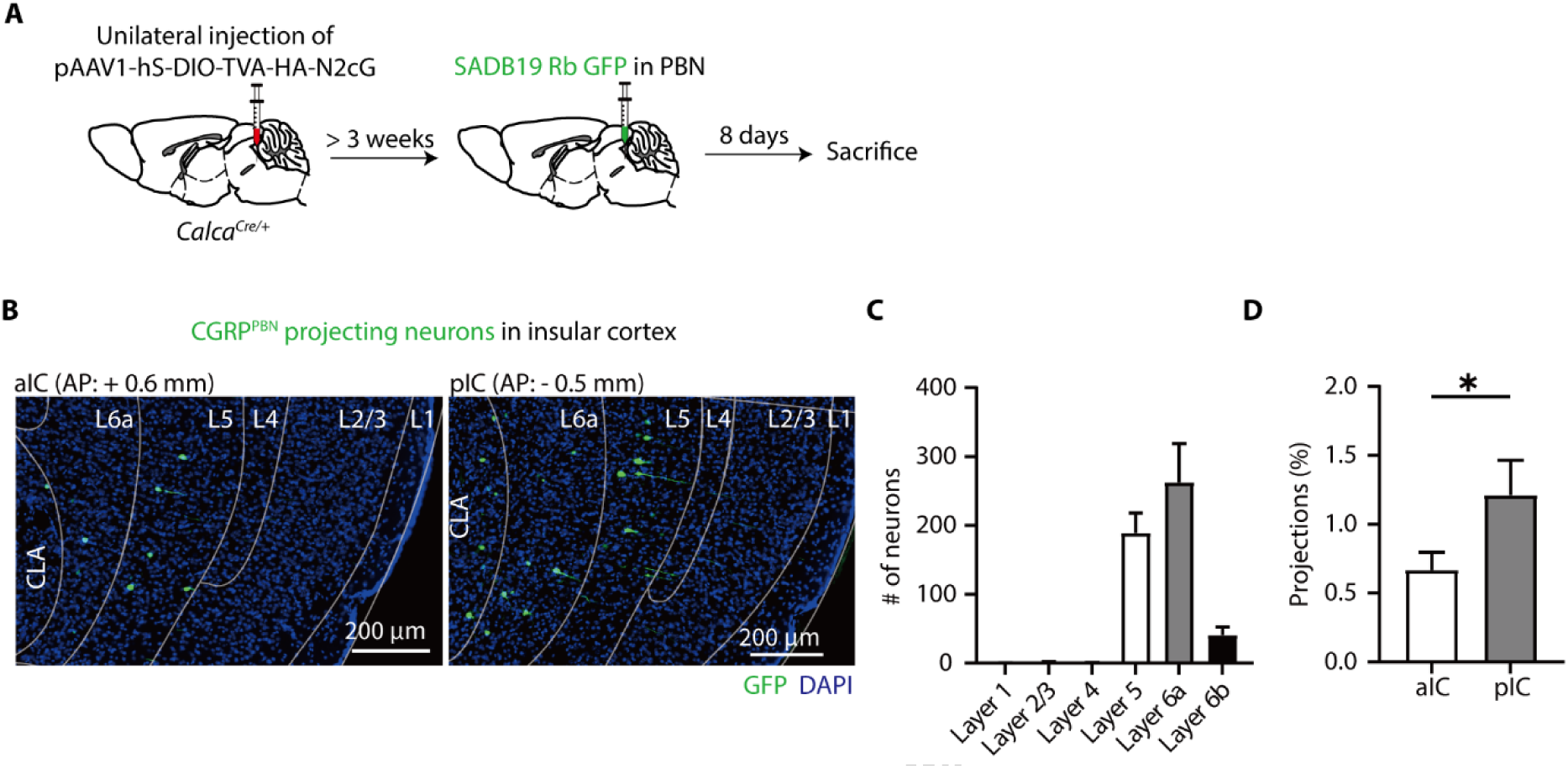
Monosynaptic retrograde tracing of CGPR^PBN^ neurons in the insular cortex (related to Figure 4). **(A)** Schematic of unilateral monosynaptic retrograde tracing of CGRP^PBN^ neurons. **(B)** Representative images of coronal brain sections showing retrogradely labeled neurons with GFP in the anterior (left) and posterior (right) parts of the insular cortex. **(C)** CGRP^PBN^-projecting posterior insular neurons are specifically located in layers 5 and 6. **(D)** Percentage of projections in the anterior and posterior insular cortex (*N*= 6 mice; Two-tailed paired t-test, t_5_= 2.815, *P*= 0.0373).

**Fig. S6.**
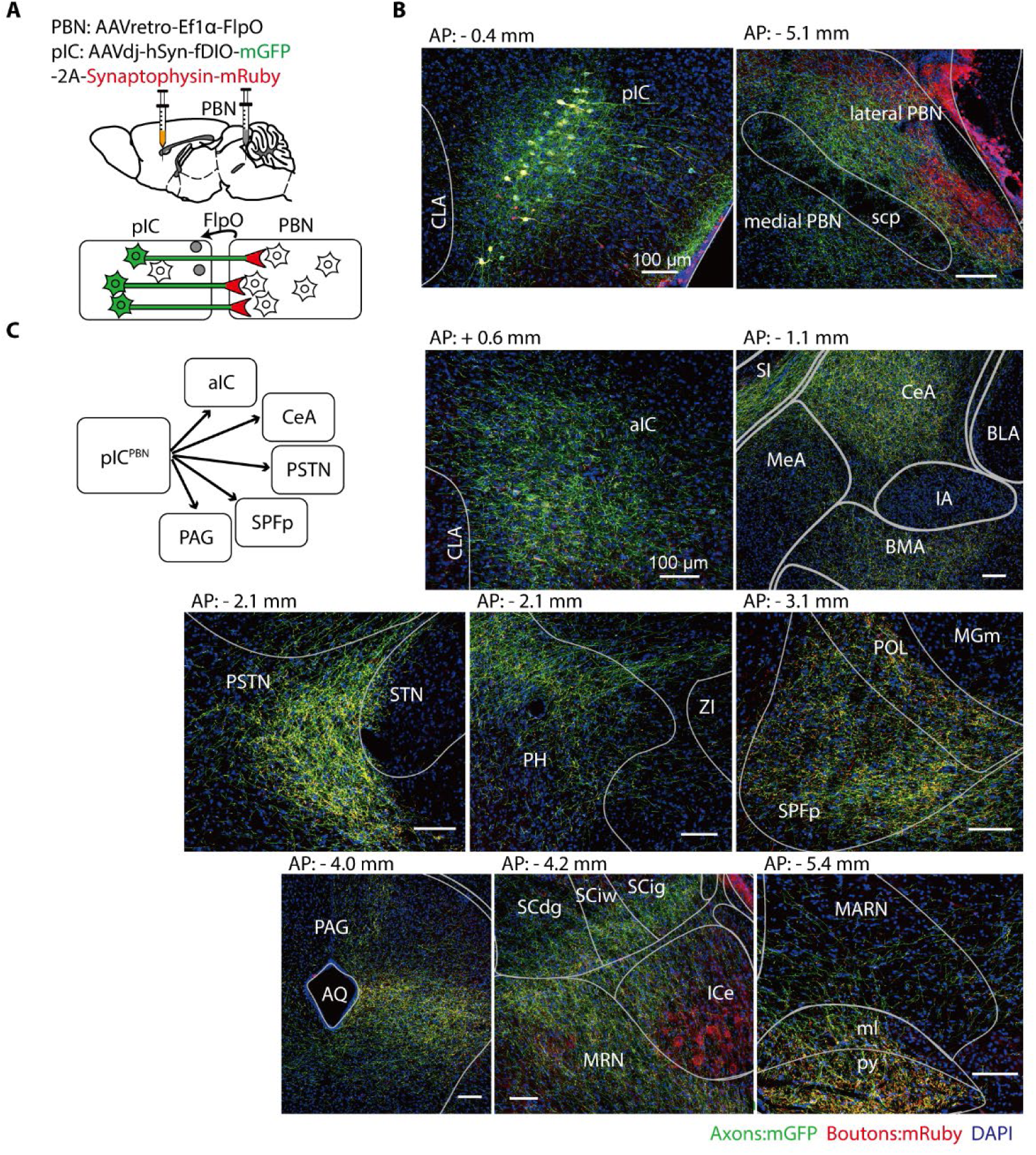
Axon tracing of pIC^→PBN^ neurons (related to Figure 6). **(A)** Schematic of unilateral axon tracing of pIC^→PBN^ neurons. Unilateral injections of AAVretro-EF1α-flpO into the PBN and AAVdj-hSyn-FLEX-mGFP-2A-Synaptophysin-mRuby into the pIC. **(B)** Representative images of pIC^→PBN^ neurons (left) and projections to the PBN (right). Scale bars, 100 µm. **(C)** Representative confocal microscopic images of coronal sections showing the axons (mGFP +) and synaptic bouton (mRuby +). Scale bars, 100 µm. **Abbreviations**: Anterior insular cortex (aIC); Basolateral amygdalar nucleus (BLA); Basomedial amygdalar nucleus (BMA); Central amygdalar nucleus (CeA); Cerebral aqueduct (AQ); Claustrum (CLA); Inferior colliculus, external nucleus (ICe); Intercalated amygdalar nucleus (IA); Magnocellular reticular nucleus (MARN); Medial amygdalar nucleus (MeA); Medial geniculate nucleus, medial part (MGm); Medial lemniscus (ml); Midbrain reticular nucleus (MRN); Parasubthalamic nucleus (PSTN); Periaqueductal gray (PAG); Posterior hypothalamic nucleus (PH); Posterior insular cortex (pIC); Posterior limiting nucleus of the thalamus (POL); Pyramid (py); Subparafascicular nucleus, parvicellular part (SPFp); Substantia innominata (SI); Subthalamic nucleus (STN); Superior cerebellar peduncle (scp); Superior colliculus, deep gray layer (SCdg); Superior colliculus, intermediate gray layer (SCig); Superior colliculus, intermediate white layer (SCiw); Zona incerta (ZI).

